# Optimizing mitochondrial maintenance in extended neuronal projections

**DOI:** 10.1101/2020.09.11.294207

**Authors:** Anamika Agrawal, Elena F. Koslover

## Abstract

Neurons rely on localized mitochondria to fulfill spatially heterogeneous metabolic demands. Mitochondrial aging occurs on timescales shorter than the neuronal lifespan, necessitating transport of fresh material from the soma. Maintaining an optimal distribution of healthy mitochondria requires an interplay between a stationary pool localized to sites of high metabolic demand and a motile pool capable of delivering new material. Interchange between these pools can occur via transient fusion / fission events or by halting and restarting entire mitochondria. Our quantitative model of neuronal mitostasis identifies key parameters that govern steady-state mitochondrial health at discrete locations. Very infrequent exchange between stationary and motile pools optimizes this system. Exchange via transient fusion allows for robust maintenance, which can be further improved by selective recycling through mitophagy. These results provide a framework for quantifying how perturbations in organelle transport and interactions affect mitochondrial homeostasis in neurons, a key aspect underlying many neurodegenerative disorders.

**Author summary:** Neurons contain long projections termed axons and dendrites and a small central body that is responsible for much of cellular biosynthesis. Mitochondria, the energy hubs of a cell, are synthesized in the soma and actively transported to distant sites of high energy demand. Given the extreme distances between these sites and the soma, maintaining distal mitochondrial health poses a substantial challenge. Defects in mitochondrial transport and maintenance are associated with several neurological disorders.

Fortunately, mitochondria stationed at distant sites can be ‘serviced’ by passing mitochondria that emerge from the soma and move around the neuron, as well as through low levels of local protein synthesis. We develop mathematical models for two strategies of mitochondrial maintenance: one with direct protein exchange between moving and stationary mitochondria (‘Space Station’) and the other with moving mitchondria occasionally replacing stationary ones at the demand sites (‘Changing of the Guard’). We find that only a few servicing events and a small motile pool form optimal conditions for maintaining mitochondrial health. The system can be improved further by selectively removing and recycling some unhealthy mitochondria. Our results are consistent with observations of mitochondrial behavior in neurons and form a basis for future quantitative study of mitochondrial maintenance.

## Introduction

Mammalian neurons, with their complex and elongated architecture, pose a unique challenge for the delivery and maintenance of cellular components. Their relatively small cell body (soma) contains the nucleus and is responsible for synthesizing all the mRNA transcripts and a large portion of the proteins delivered to distal regions [1, 2]. For some proteins, local translation at distal outposts has been shown to play an important role in maintaining homeostasis [3–5]. However, this approach still requires the long-range delivery of mRNA from the nucleus [6–8]. Neurons thus rely on packaging components into vesicular organelles or RNA-protein granules, which are actively transported by molecular motors moving along microtubule tracks to the most distant regions of the cell. Such long-distance transport is critical for neuronal growth and repair [9–11], synapse formation and function [12–14], and synaptic plasticity [15, 16]. Furthermore, transport is required to maintain steady-state homeostasis of protein and mRNA levels. While neurons can have a lifespan of decades, the turnover of active mRNA is thought to happen on timescales of hours [17], and proteins are degraded over the course of a few days [18]. Efficient, perpetual delivery through transport from the cell body is thus required to replenish degraded components and maintain neuronal function.

A particular challenge for cellular homeostasis is supplying the spatially heterogeneous metabolic needs of neuronal projections. These projections, which stretch up to several hundred microns for dendrites and over a meter for axons, contain localized structures with elevated metabolic demand. For example, distal axonal structures such as presynaptic terminals and growth cones are known to have especially high ATP needs [19, 20]. In myelinated axons, the nodes of Ranvier can be separated by hundreds of microns [21] and contain high concentrations of ion pumps that are thought to correspond to locally elevated metabolic demand [22]. Postsynaptic dendritic spines also have high metabolic needs to support both ion pumping and local translation needed for proteostasis and plasticity [23, 24].

Reliable ATP production at sites of high metabolic demand is believed to rely on the localization of nearby mitochondria. These organelles serve as ATP-generating powerhouses and also help maintain Ca^+2^ homeostasis in the cytoplasm. Locally stationed mitochondria are a key source of fuel to power activity-dependent protein synthesis in dendritic compartments [23]. Mitochondria are also known to accumulate in presynaptic terminals [25] and in juxtaparanodal regions next to nodes of Ranvier in myelinated axons [26, 27]. Mitochondria are manufactured primarily in the soma [28] and delivered throughout neuronal projections via motor-driven transport [29, 30]. Delivery of mitochondria to specific regions relies on localized signals that trigger halting and switching into a stationary state. Several mechanisms for stopping motile mitochondria have been identified [30], including Ca^+2^-dependent immobilization [31], glucose-dependent arrest [32], and syntaphillin-based anchoring [33]. Interestingly, 60-90% of axonal mitochondria are observed to be stationary, while the remainder are roughly equally split between an anterograde and retrograde motile population [30].

Mitochondria are one of many neuronal components whose delivery relies on local stop signals rather than a global addressing system. Dense core vesicles carrying neuropeptides [34] and RNA-protein particles [15] are distributed by sporadic capture at sequential synaptic sites as they circulate through the dendrites. Such delivery systems have been described by a “sushi belt model”, where local sites trigger removal of components from an effective constantly moving conveyor belt [15, 35]. Quantitative analysis of the sushi belt model indicates that these systems face severe trade-offs between speed and accuracy, so that frequent halting interferes with the ability of transported components to reach the most distal regions [35]. The efficiency of a sushi-belt delivery system for steady-state replenishment of degrading components has not previously been addressed, and forms a key aspect of the current work.

The placement of stationary mitochondria at distal regions raises a challenging problem of “mitostasis” (mitochondrial homeostasis) [29]. Mitochondria are known to age over time, decreasing their membrane potential and ATP production efficiency [36], possibly as a result of damage caused by reactive oxygen species [37]. Defects in mitochondrial transport and quality control are associated with a variety of neurological diseases, including Alzheimer’s, Huntington’s, and Parkinson’s disease [38, 39]. Individual mitochondrial proteins have typical half-lives in the range of 2-14 days [18, 40]. Because most mitochondrial proteins are encoded in the nucleus, this implies that a distally stationed mitochondrion will experience significant degradation of its protein complement over the days to weeks time-scale. While local translation may replenish some of this protein content, not all mRNAs are found outside the soma [41], and the protein synthesis capacity of the axon is only a small fraction of that in the cell body [2]. Maintenance of a healthy population of mitochondria thus requires the continual delivery of new mitochondrial material to distal outposts. While some of this delivery could be accomplished by vesicular transport followed by local import, most nuclear-derived mitochondrial content is believed to be transported by a motile population of mitochondria themselves [29, 30].

Two main qualitative models have been proposed for the replenishment of aging neuronal mitochondria through long-range transport [29]. One, termed the ‘Changing-of-the-Guard’ (CoG) model, relies on individual mitochondria switching between the stationary and the motile pools. This enables newly synthesized mitochondria moving anterograde from the cell soma to halt at regions of high metabolic demand, while stationary mitochondria begin moving again to reach other regions or return to the soma for recycling. An alternative approach is the ‘Space Station’ (SS) model, in which a pool of mitochondria remains permanently stationed at distal sites. In this model, new protein components are delivered by passing motile mitochondria that undergo transient fusion and fission events with the stationary organelles.

Fusion and fission dynamics enable the exchange of membrane and matrix contents between distinct mitochondria [42–44]. In certain cell types, fusion allows the formation of extended mitochondrial networks that have been hypothesized to contribute to the efficient mixing of components that determine mitochondrial health [45, 46]. In neuronal axons, extensive fusion into a mitochondrial network is not observed but transient kiss-and-run events, consisting of a rapid fusion and fission cycle between two passing mitochondria, can nevertheless allow for component exchange [47, 48].

Neuronal mitostasis is also facilitated by a recycling pathway termed “mitophagy” [49]. In this pathway, unhealthy mitochondria with low membrane potential are marked by an accumulation of protein kinase PINK1 which recruits ubiquitin ligase Parkin, leading to ubiquitination of the mitochondrial surface [30, 49]. Autophagosomes (originating primarily at distal cell tips) move in the retrograde direction, engulf tagged damaged mitochondria, and carry them back to the cell body for recycling [29, 50].

Past theoretical studies of mitochondrial maintenance have focused on systems with extensive mitochondrial fusion and asymmetric fission [51, 52]. When mitochondrial health is determined by discrete factors in low copy numbers, fission events can stochastically result in particularly unhealthy daughter mitochondria that can then be targeted for degradation through mitophagy. The combination of discrete health units and selective autophagy has been shown to be sufficient for maintaining a healthy mitochondrial population [51, 52]. Recent studies have also begun to explore the role of spatial distribution, with randomly directed active transport and proximity-dependent fusion [52]. However, no quantitative model has yet attempted to address mitochondrial maintenance in the uniquely extended geometry of neuronal projections. This cellular system relies on long-range mitochondrial transport and faces the challenge of positioning mitochondria at specific distal regions of high metabolic demand.

In this work, we develop quantitative models for the coupling between mitochondrial transport and mitochondrial homeostasis in neuronal projections. We treat mitochondrial health as a continuously decaying parameter along the axon and assume that the generation of some of the critical mitochondrial material essential for health is limited to the cell body. Our models encompass both the ‘Space Shuttle’ and the ‘Changing-of-the-Guard’ qualitative mechanisms for mitochondrial exchange. In contrast to past work [52], we focus not only on total mitochondrial health, but also on the distribution of healthy mitochondria among localized distal regions of high metabolic demand. The balance between mitochondrial transport and localization, in the face of decaying mitochondrial health, serves as a bridge between spatially-resolved models of neuronal distribution [34, 35] and global models of mitochondrial maintenance [51, 52].

We leverage both analytically tractable mean-field methods and discrete stochastic simulations to explore the equivalence between the SS and CoG mechanisms, delineate the parameter values that maximize mitochondrial health, and establish the existence of an optimum health threshold for mitophagy. This modeling effort fills a gap in our existing understanding of how neurons accomodate the trade-offs inherent in maintaining mitochondrial homeostasis while positioning mitochondria in regions far distant from the primary site of protein and organelle biogenesis.

## Models for mitochondrial maintenance

Two inherent constraints pose a challenge for maintaining distal mitochondrial outposts in a linearly extended region such as mammalian axons. First, mitochondrial health is assumed to be dependent on components (*eg:* proteins) that degrade rapidly compared to the cell lifespan and must be manufactured primarily in the cell soma at one end of the domain. The limitation of long-range transport from the soma can, in part, be bypassed by local protein translation, a phenomenon whose role has been increasingly appreciated in recent years [3–5]. Local protein synthesis can both maintain distal levels of fast-degrading proteins and allow for rapid response to changing local conditions [5, 54]. It should be noted, however, that local translation shifts the burden of transport and maintenance to mRNA, which tends to have even shorter life-times than many proteins [55–57]. We briefly explore the contribution of local translation to our model in a subsequent section, but focus primarily on mitochondrial health components that must be transported from the cell body.

A second key assumption is the existence of discrete “demand sites” interspersed throughout the domain, which have a particularly high need for a healthy population of localized mitochondria. For simplicity, we treat the demand sites as very narrow slices of the domain and focus on mitochondrial health specifically within those slices – an extreme form of spatial heterogeneity in metabolic demand. In the opposite extreme of wholly homogeneous metabolic demand, the optimum maintenance strategy becomes simple: all mitochondria should move back and forth through the cell as rapidly as possible (within the limits of other constraints such as energetic requirements of transport) in order to ensure frequent replenishment of proteins at the cell soma.

In a system with discrete demand sites, optimizing metabolism requires the establishment of a population of stationary mitochondria specifically at those sites. However, this stationary population must be continuously replenished by a flux of either new proteins or entirely new mitochondria. We develop two quantitative models for mitochondrial replenishment (the SS and CoG model, illustrated in Fig. 1), in keeping with recent qualitative proposals for mitochondrial maintenance mechanisms [29].

**Fig 1.**
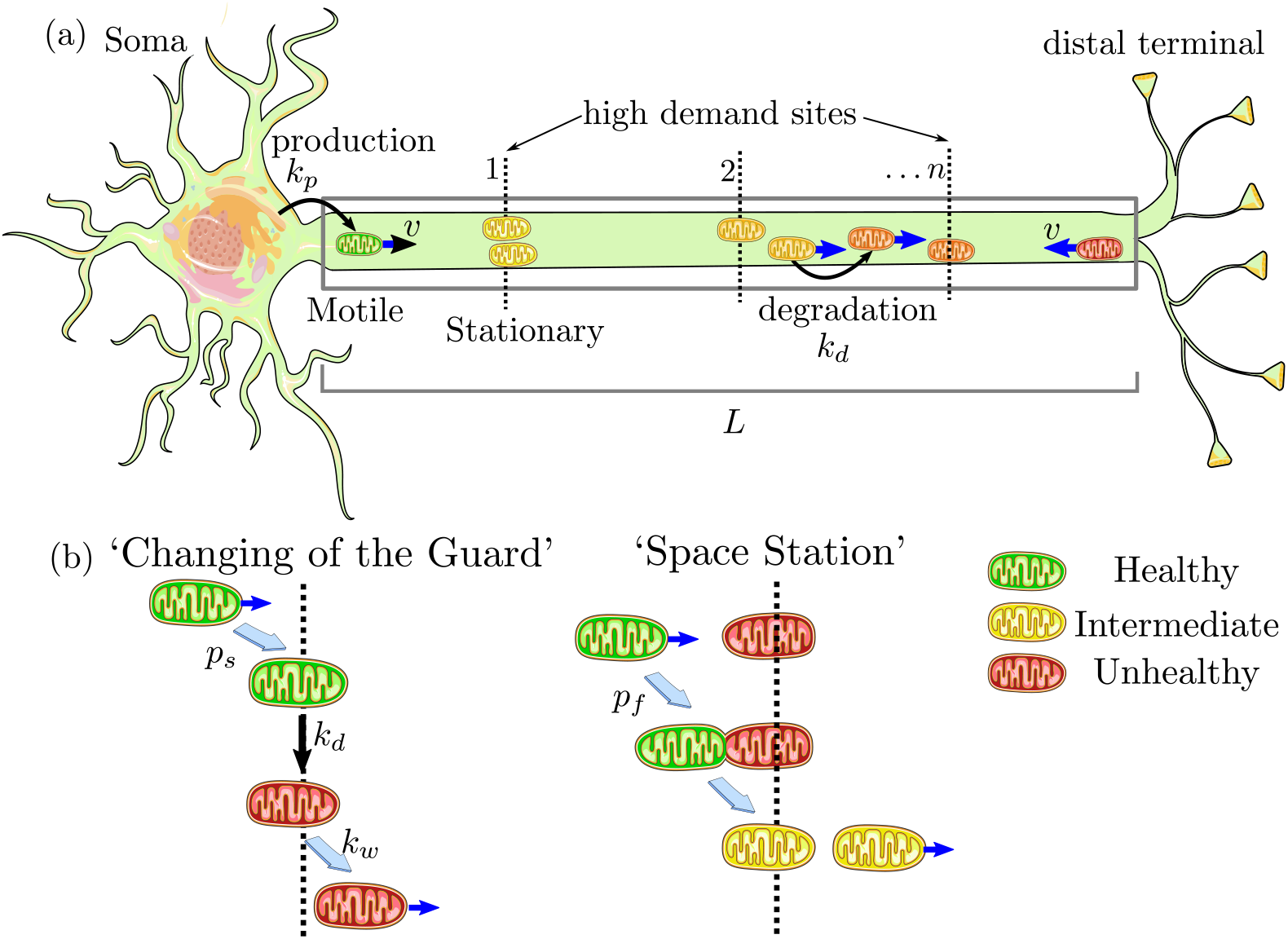
Schematic of quantitative models for mitochondrial maintenance in long neuronal projections. (a) Mitochondria are produced at the soma at rate *k*_*p*_, move processively with velocity *v*, and can stop at one of *n* discrete sites with high metabolic demand. Mitochondrial health degrades continuously with rate *k*_*d*_. Gray box represents the modeled linear domain of length *L*. (b) Two models for mitochondrial exchange at demand sites. In the CoG model, stationary mitochondria re-enter the motile population with rate *k*_*w*_ while passing motile mitochondria stop with probability *p*_*s*_. Stopping and restarting rates are independent of mitochondrial health; on average, however, restarting mitochondria will be less healthy than when they first stopped at the site, due to degradation in the stationary state. In the SS model, transient fusion events occur with probability *p*_*f*_ each time a motile mitochondrion passes a permanently stationary one [53].

In both cases, we assume that there are *n* discrete point-like demand sites at positions *x*_*i*_, placed at equal separations over a linear domain of length *L* (Fig. 1a). This long linear domain mimics the extended axonal regions imaged in cell cultures [32] and serves as a simplification that enables us to focus on the interplay of dynamic processes for determining mitochondrial homeostasis. The effect of more complicated branched geometries observed *in vivo* is explored briefly in Supplementary Material.

We treat individual mitochondrial health as a continuous quantity that is set to 1 when the mitochondrion leaves the cell body and decays exponentially with constant rate *k*_*d*_ as it travels down the axon. This health can be thought of as proportional to the copy number of an unspecified critical mitochondrial protein that is manufactured in the soma and is turned over during the lifetime of the mitochondrion. Other studies have also focused on the membrane potential and the accumulation of mutations in mitochondrial DNA as markers for mitochondrial health [51, 52]. With an appropriate change of units, the mitochondrial health in our model can equivalently represent any of these quantities, requiring only that they change in a continuous manner, with a single well-defined decay rate. Although a variety of factors may contribute to mitochondrial aging [58], we include only one decay rate for a single limiting health factor in our simplified model. More complicated systems that incorporate multiple decaying components or different decay rates for the more metabolically active stationary mitochondria versus motile mitochondria are left for future expansions of the basic model. For clarity of exposition, we will assume that health corresponds to protein content in the remainder of the discussion.

The metabolic health at each demand site (*H*_*i*_) is given by the total protein content for all mitochondria stationed at the site. Since each site is assumed to be very narrow (on the order of a few microns in an axonal length of mm to cm), the motile mitochondria spend negligible time at these sites and can be neglected from our calculations of demand site health. We consider two utility functions for overall metabolic health. The first is 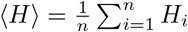, the average health over all demand sites. The second is the health level at the most distal demand region (*H*_*n*_), which serves as the lowest value across all the sites. Optimizing ⟨*H*⟩ requires maximizing the sum total metabolic rate in many regions of interest. Such an optimum could correspond to an uneven distribution of mitochondrial health between individual regions, with the distal demand sites being poorly supplied, while the proximal sites maintain a high health level. By contrast, optimizing *H*_*n*_ ensures that each individual region exhibits, to the greatest extent possible, a high level of mitochondrial health. Increasing the total number of mitochondria in the domain (*M*) will predictably raise both metrics. At the same time, requiring a fixed total number of mitochondria to service a greater number of demand sites should decrease the average health at each site. Consequently, we normalize the demand-site health levels by the total number of mitochondria per site:

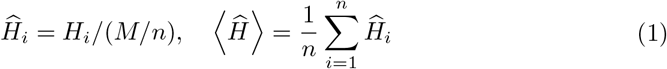

An individual mitochondrion can either be motile (moving with velocity ±*v*) or stationary (at one of the demand sites). For simplicity, we neglect transient pausing of motile mitochondria and treat them as having a constant effective velocity outside of the demand regions. In keeping with experimental observations of axonal mitochondria [59], a motile mitochondrion is assumed to move processively in the anterograde or retrograde direction, with no switching of directionality in the bulk of the domain. This assumption is relaxed in the Changing-of-the-Guard model, where stationary mitochondria can restart in either the anterograde or retrograde direction. The fate of mitochondria when they reach the terminus of an axon may involve some combination of local degradation [60] and recirculation towards the soma [61–63]. Given that the retrograde and anterograge flux of mitochondria are similar in axons [32, 62, 64], we assume that most of the organelles return to the cell body for recycling. Modeled mitochondria that reach the end of the domain (length *L*) are thus assumed to instantaneously switch to retrograde motion. A retrograde mitochondrion that returns to the soma at *x* = 0 is removed from the domain. To maintain a steady-state total mitochondrial number, anterograde mitochondria with full health are produced at the somal end with rate *k*_*p*_.

The difference between the two models lies in the ability of individual mitochondria to switch between stationary and motile pools or to exchange protein content with other mitochondria, as described below.

### Changing-of-the-Guard (CoG) model

One mechanism for maintaining a healthy population of localized mitochondria is by allowing occasional interchange between the pool of motile mitochondria and those stationed at demand regions. In the absence of fusion and protein exchange, motile mitochondria are generally younger and healthier than stationary ones, and the latter must be removed and recycled sufficiently quickly to maintain overall mitochondrial health at the regions of interest.

The simplest version of this model assumes that the rate of restarting stationary mitochondria is independent of their health or demand site position along the axon. An expanded version that incorporates mitophagy to selectively remove unhealthy mitochondria is considered in a subsequent section. Here, we assume each moving mitochondrion has a constant probability *p*_*s*_ for switching to the stationary state each time it passes a demand region. A stationary mitochondrion restarts motility with fixed rate constant *k*_*w*_. Upon re-entering the motile state, it is equally likely to move in either the anterograde or retrograde direction. Stochastic agent-based simulations of this model are illustrated in Fig. 2a and Supplemental Video S1.

**Fig 2.**
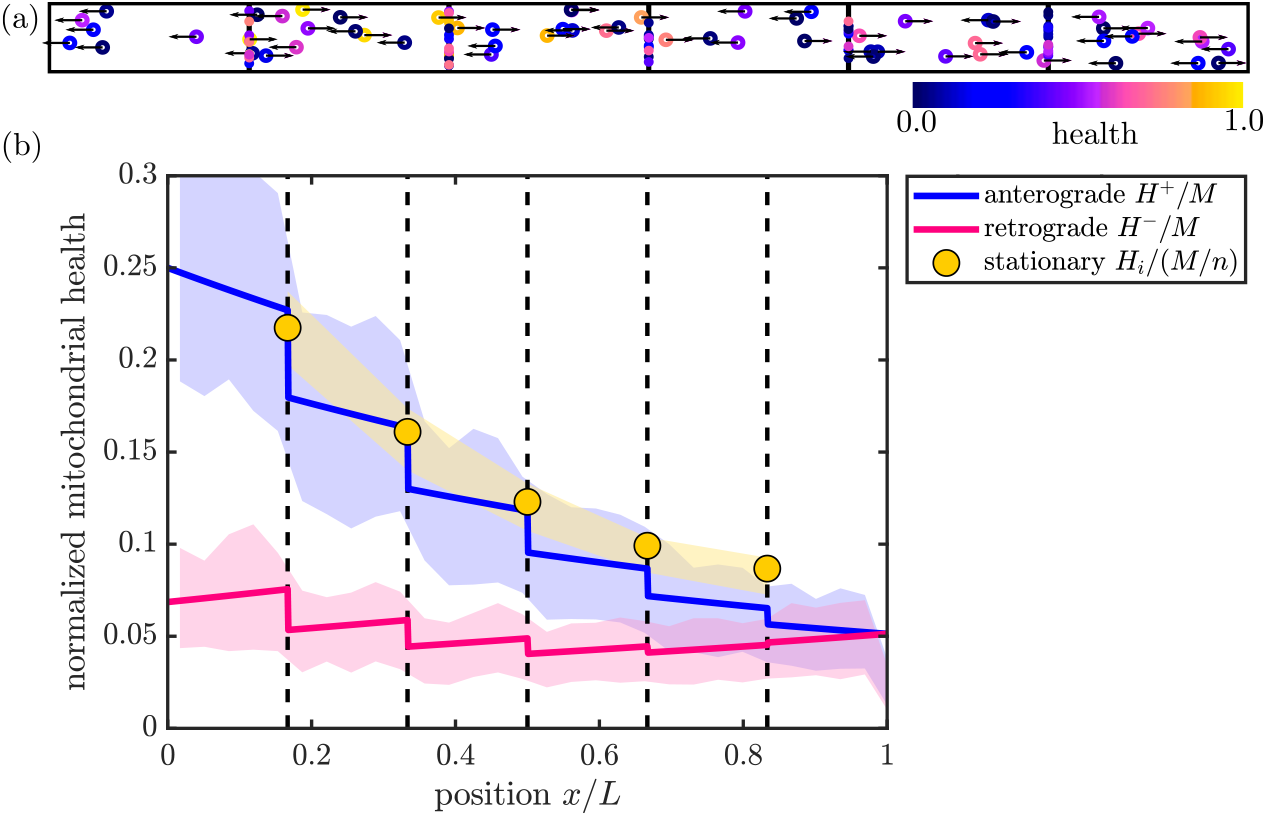
Comparison of mean-field and discrete stochastic models. (a) Snapshot of stochastic simulation of the CoG model, with *M* = 100. (b) Steady-state solution for mitochondrial health in the CoG model (Eq. 2). Solid curves show linear density of mitochondrial health in anterograde (blue) and retrograde (magenta) mitochondria, normalized by total number of mitochondria in the domain. Yellow circles show total health at each of the discrete demand sites (dashed black lines), normalized by the total number of mitochondria per region. Shaded regions show corresponding quantities from discrete stochastic simulations (mean ± standard deviation) with *M* = 1500 mitochondria in the domain. Parameters used in (a) and (b): *n* = 5, *p*_*s*_ = 0.4, *f*_*s*_ = 0.5, *k*_*d*_*L/v* = 0.6. Corresponding results for the SS model are provided in Supporting Information (S4 Fig).

A mean-field description of this model tracks the behavior of *H*^*±*^(*x, t*) (the linear density of mitochondrial health moving in the anterograde (*H*^+^) and retrograde (*H*^*−*^) directions) and *H*_*i*_ (the total health of stationary mitochondria at demand site *i*). These quantitites evolve according to the following set of equations:

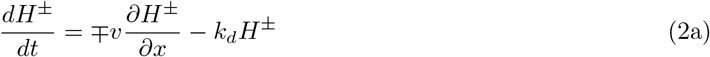

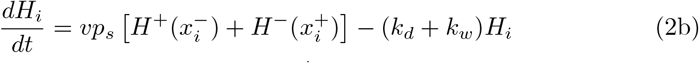

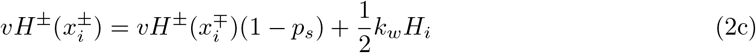

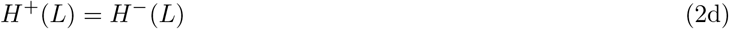

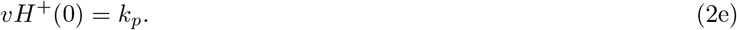

Equation 2a describes transport and decay of health (or protein content) in motile mitochondria. The distributions *H*^*±*^(*x, t*) are continuous on each interval (*x*_*i*_, *x*_*i*+1_), with discontinuities at the demand sites. Equation 2b encompasses switching between the stationary and motile populations, with 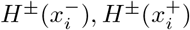 referring to the limit of the distribution approaching the *i*^th^ site from the negative and from the positive side, respectively. Equation 2c gives the boundary condition at each demand site, where the flux of outgoing proteins from one side of the site must equal the flux of incoming proteins that pass without stopping from the other side, as well as the flux of stopped proteins that re-enter the motile state in that direction. Equation 2d describes switching of anterograde to retrograde motile mitochondria at the distal end of the domain. Finally, Eq. 2e gives the production rate of healthy mitochondria at the proximal end of the domain, assuming a health level of 1 for each newly-made mitochondrion.

The distribution of mitochondria themselves obeys the same set of equations (Eq. 2) with the *k*_*d*_ terms removed. We define *ρ* as the steady-state density of all motile mitochondria and *S* as the steady-state number of stationary mitochondria at each demand site. At steady state, *ρ* is constant throughout the domain and *S* is the same in all demand sites, with

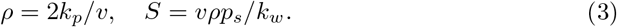

The total number of mitochondria in the domain is given by *M* = *ρL* + *nS*. We use the total mitochondria *M* as a control parameter in our models, allowing *k*_*p*_ to vary as needed in order to maintain a given value of *M*. This approach represents a system where the total mitochondrial content in the domain is limited, so that increasing the number of stationary mitochondria comes at the expense of having a smaller motile population. The steady-state fraction of mitochondria in the stationary pool is defined as *f*_*s*_ and can be expressed as

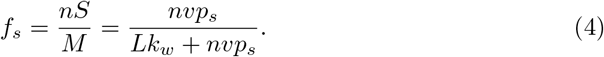

The quantity *f*_*s*_ encompasses the trade-off between health replenishment by motile mitochondria and metabolic servicing of localized sites by stationary mitochondria, making it a convenient control parameter to explore this balance throughout the rest of our study.

At steady state, Eq. 2 can be solved analytically to give the distribution of mitochondrial health throughout the domain and at each demand site (Fig. 2b; see Materials and Methods for details). The mean-field model accurately describes the steady-state distributions when averaged over many realizations of stochastic trajectories for discrete mitochondria (Fig. 2b).

In each of the regions between demand sites, the steady-state anterograde protein density drops exponentially as 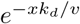 with distance from the soma. The reverse relationship holds for retrograde-moving protein density, which decreases exponentially towards the soma. It should be noted that the overall mitochondrial health decreases with increasing distance from the soma – a fundamental consequence of long-range transport and continual protein turnover. Interestingly, a modest correlation between mitochondrial aging and distance from the nucleus has been experimentally observed in hippocampal neurites [65]. Furthermore, some measurements indicate that retrograde moving mitochondria have lower health than anterograde ones [66] as also seen in Fig. 2.

### Space Station (SS) model

An alternate approach to mitochondrial homeostasis relies on maintaning a pool of permanently stationed mitochondria in each demand region, whose contents can be replenished by transient fusion and fission with the motile population. Such transient ‘kiss-and-run’ events have been observed to allow for exchange of mitochondrial membrane and matrix proteins in H9c2 and INS-1 cells [47].

In our quantitative Space Station model, permanently stationary mitochondria are placed at the high demand sites, with *S* = *f*_*s*_*M/n* mitochondria per site. We assume the mitochondria maintain their discrete identities following a transient fusion/fission event, and that the protein content of the two mitochondria is fully equilibrated in each such event. Every time a motile mitochondrion passes a demand site, it has the opportunity to fuse with each of the stationary mitochondria present at that site, in sequence. Each transient fusion/fission event is successful with probability *p*_*f*_, and the choice to fuse with each stationary mitochondrion is independent of health levels or prior fusion events. Following an instantaneous fusion/fission cycle, the health levels of the mitochondria are equilibrated, so that a protein in a motile mitochondrion has probability *p*_*f*_ */*2 of switching to a stationary one each time there is a passage event. The rate at which a stationary protein re-enters a motile state in this model is given by *p*_*f*_ */*2 times the flux of motile mitochondria passing the stationary organelle containing that protein. At steady state, this gives an effective restarting rate of

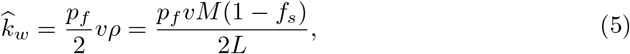

where *ρ* is the linear density of motile mitochondria and *f*_*s*_ is the fraction of mitochondria in the permanent stationary state.

The mean-field equations for the SS model are derived in the Methods section. For the case with only one mitochondrion per site (*S* = 1), they are mathematically equivalent to the CoG model, with the variable replacements *p*_*s*_ → *p*_*f*_ */*2 and 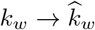.

Stochastic simulations of the SS model with discrete mitochondria are illustrated in Supplemental Video S2, and the resulting steady-state mitochondrial health matches well to the mean-field model (S4 Fig).

### Model comparison

In order to compare the efficiency of mitochondrial maintenance via the CoG and SS models, we must first consider the different control parameters involved. Parameter definitions for both models are summarized in the Methods section. The parameters *k*_*d*_ (protein decay rate), *M* (total mitochondria), *N* (number of demand sites), *v* (motile mitochondria speed), and *L* (domain length) play the same role in both models. The CoG model is further parameterized by *k*_*w*_ (restarting rate for a stationary mitochondrion) and *p*_*s*_ (stopping probability for motile mitochondrion at each demand site). The SS model, on the other hand, has the parameters *S* (fixed number of stationary mitochondria per site) and *p*_*f*_ (probability of fusion with each stationary mitochondrion). These last two parameters define two key features of the maintenance mechanism: the steady-state fraction of mitochondria in the stationary pool (*f*_*s*_), and the probability that a protein within a motile mitochondria will transition into a stationary state while passing each demand site 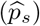. If these two control parameters are fixed, then for the CoG model we have 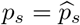 and *f*_*s*_ given by Eq. 4. For the SS model, we set *f*_*s*_ = *nS/M* directly. Because a motile mitochondrion has the opportunity to fuse sequentially with multiple stationary organelles at each demand site, the overall probability that a protein will transition to a stationary state at that site is given by

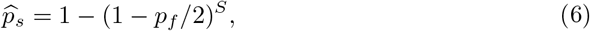

allowing a corresponding *p*_*f*_ value to be set for each effective stopping probability 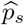.

When there is just one mitochondrion stationed in each demand site (*S* = 1), the two mean-field models become mathematically identical for fixed values of *f*_*s*_ and 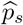. In general, the two models give very similar results so long as the stopping probability at each demand site is small 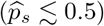, as shown in Fig. 3. In this limit, 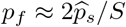 and 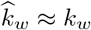 (from Eq. 4–5). For large values of 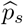, the SS model is seen to yield slightly lower mitochondrial health than the CoG model (Fig. 3). It should be noted that for a fixed value of *f*_*s*_, the SS model can only reach a limited maximum value of the effective stopping probability 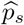. In fact, even if fusion events occur with each stationary mitochondrion (*p*_*f*_ = 1), a motile mitochondrial component always has a non-zero chance of continuing in its motile carrier rather than being left behind at the demand site. In the remainder of this work, we focus primarily on low values of 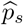, a regime where the two models give essentially equivalent mean-field results.

**Fig 3.**
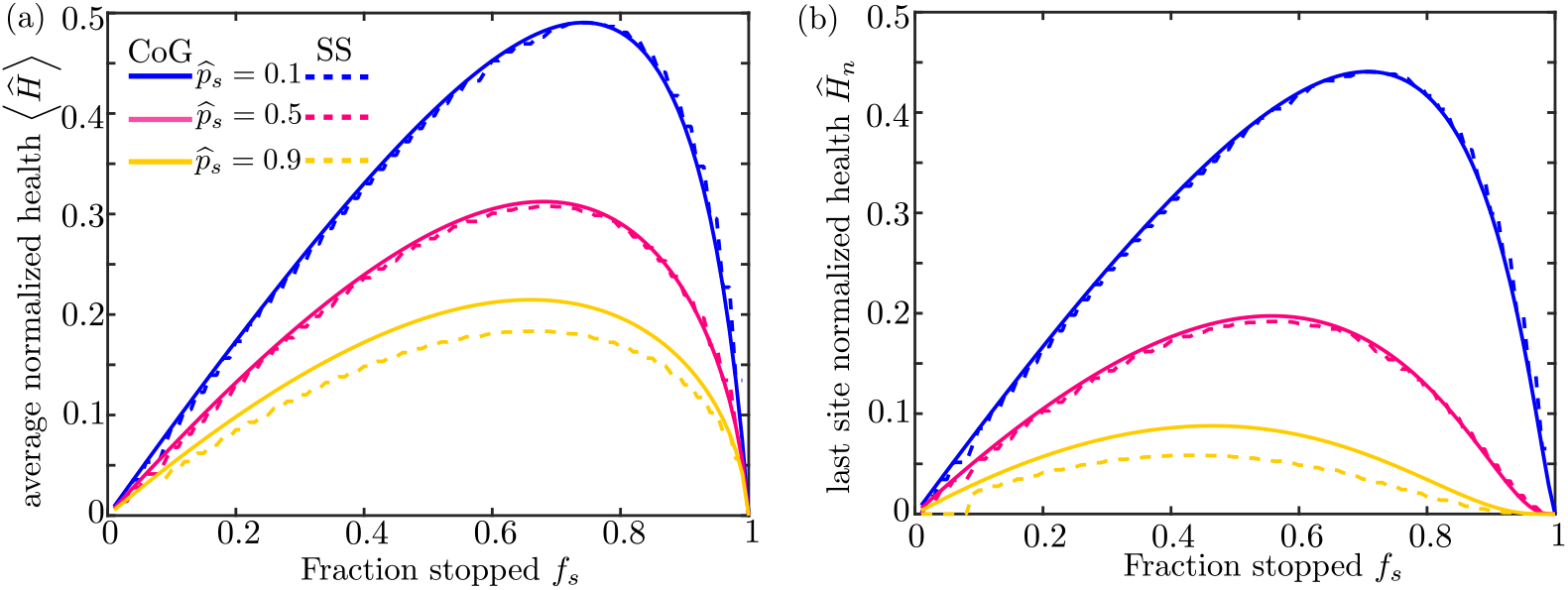
Comparison of mitochondrial maintenance models for matched parameter values. (a) Steady-state normalized mitochondrial health averaged over all demand regions is computed with the CoG model (solid lines) and the SS model (dashed lines) as a function of the fraction of stopped mitochondria (*f*_*s*_) for three different values of the effective protein stopping probability 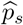. (b) Corresponding plots of the normalized health in the most distal demand site, for both models. All values are computed with *M* = 1500, 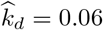, and *n* = 30.

To reduce the number of relevant model parameters, we can apply dimensional analysis, non-dimensionalizing all length units by the domain length *L*, and time units by *L/v* (the time for a motile mitochondrion to traverse the domain). The remaining dimensionless parameters are the non-dimensionalized decay rate 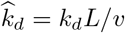, the number of mitochondria in the domain *M*, the number of demand sites *n*, the fraction of mitochondria in the stationary population *f*_*s*_, and the effective protein stopping probability at each demand site 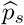.

In the CoG model, the total number of mitochondria is directly proportional to the production rate *k*_*p*_, which sets the boundary condition at the proximal edge of the domain (Eq. 2e). Consequently, the normalized health values ⟨ *Ĥ* ⟩ and *Ĥ*_*n*_ (which are scaled by the number of mitochondria per site) are independent of the total number of mitochondria in the system. The same relationship is seen for the SS model (S5 Fig). The chosen normalization embodies the assumption that the total amount of mitochondrial material in the cell is limited by other constraints, and that we are interested in modeling the overall mitochondrial health for a particular fixed number of total mitochondria.

In both maintenance models, mitochondrial proteins are removed from the system via two mechanisms: they undergo decay (with rate 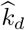) or they are recycled in the soma when a motile mitochondrion returns to its starting point. The total protein content in the system therefore depends on the time-scale for a given protein to return to the soma, and hence on the number of stops made by a given protein in the domain. The average total number of stops during a protein’s back and forth journey through the cell is given by 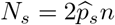. As shown in Figure 4, it is this parameter rather than the specific number of regions *n* and stopping probability *p*_*s*_ that primarily determines the mitochondrial health at each demand site.

**Fig 4.**
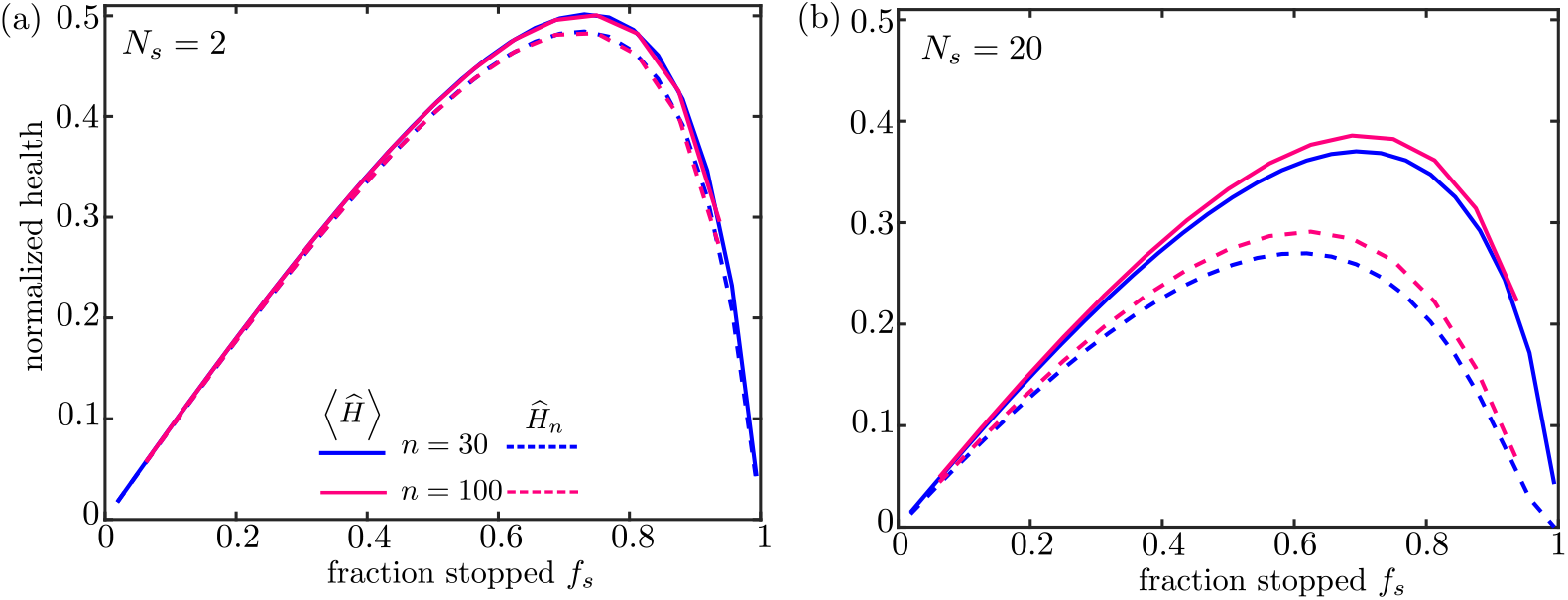
Mitochondrial health as a function of key dimensionless parameters. Solid curves show normalized average health over all demand sites; dashed curves show normalized health at the most distal site. The number of demand sites is set to *n* = 30 (blue) or *n* = 100 (magenta). For each fraction of stationary mitochondria the fusion probability is adjusted to give a fixed number of stopping events for an individual protein traversing the domain: (a) *N*_*s*_ = 2 and (b) *N*_*s*_ = 20. All values shown are for the SS model, with *M* = 1600 and 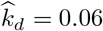. Results for different values of *M* are provided in Supporting Information (S5 Fig).

For the maintenance models proposed here, there are thus three primary parameters of relevance: the steady-state fraction of stationary mitochondria *f*_*s*_, the number of stops expected for an individual protein while moving along the domain *N*_*s*_, and the dimensionless protein decay rate 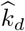. The fraction *f*_*s*_ embodies the trade-off between mitochondrial transport and localization at distal sites. The number of stops *N*_*s*_ quantifies how often proteins exchange between the mobile and stationary populations. The dimensionless decay rate 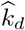 indicates how much health decay is expected for a mitochondrion that moves without stopping down an entire axon. Its value depends on the protein lifetime (a few days for mitochondrial proteins [18, 40]), the average speed of motile mitochondria (∼ 0.5*µ*m/s [59]), and the length of the axon itself. Axon lengths can vary widely, from hundreds of microns up to the meter-long axons in the human sciatic nerve [29]. Because mitochondrial maintenance is particularly challenging when mitochondria must be stationed far away from the cell nucleus, we focus here on long axons in the range of 1 − 10 cm. This range corresponds approximately to the average axon lengths measured in callossal neurons in monkey and human brains [67]. Estimating *k*_*d*_ ≈ (4days)^*−*1^ and *v* ≈ 0.5*µ*m/s, this gives dimensionless decay rates of 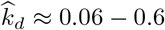.

In Supplementary Material, we include an expansion of the SS model to a branched tree geometry, rather than linear. Comparing steady-state mitochondrial health for matched values of 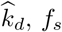, and *N*_*s*_, gives relatively similar results for the branched and linear geometries (S2 Fig), despite the exponentially greater volume, and hence much lower overall mitochondrial density, in the branched structure. This result further highlights the importance of the three main parameters identified here in determining mitochondrial health.

## Optimizing mitochondrial maintenance

We consider the average health of the system as a function of the mitochondrial transport parameters *f*_*s*_ and *N*_*s*_ (Fig. 5). An optimal value of the average health is observed at intermediate values of both the stopped fraction and the number of stopping events. When too few mitochondria are in the stationary state (low *f*_*s*_), then the total health at the demand regions is low simply because there are very few mitochondria present in those narrow regions. On the other hand, if too many mitochondria are stationary (high *f*_*s*_), the pool of motile mitochondria available for replenishing decayed proteins at the demand sites becomes very sparse and the health of the system diminishes. The optimum with respect to *f*_*s*_ relies on the assumption that the total mitochondrial content in the domain is limited, embodying a trade-off between the stationary and motile populations.

**Fig 5.**
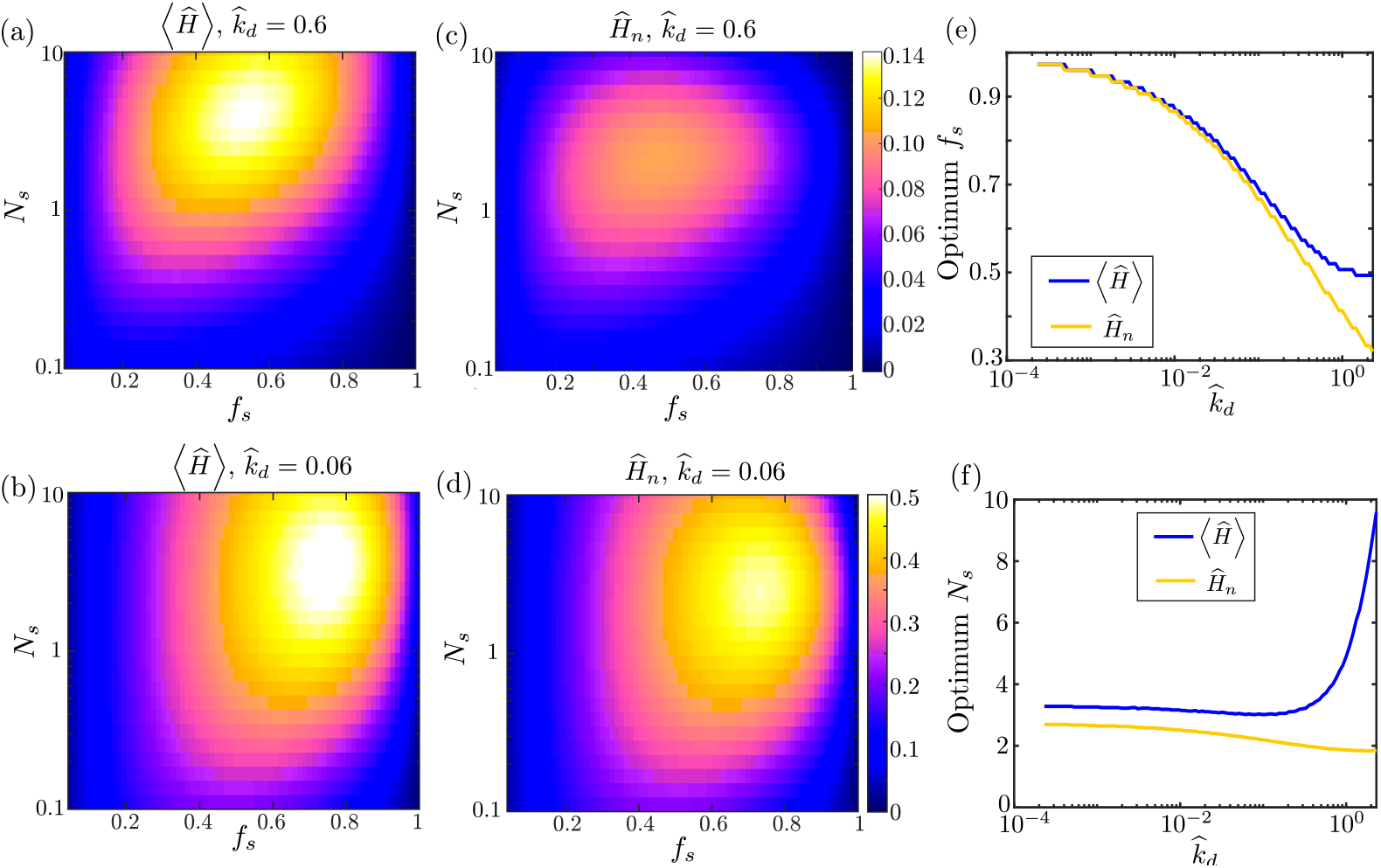
Optimizing mitochondrial health through variation of transport parameters. (a-b) Average health across all demand regions as a function of fraction of stopped mitochondria (*f*_*s*_) and number of stopping events (*N*_*s*_), for two dimensionless decay rates 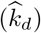. (c-d) Mitochondrial health at the most distal demand site, for two different decay rates. (e-f) Values of the *f*_*s*_ and *N*_*s*_ parameters that correspond to maximum average health (blue curves) or last region health (yellow curves). Optimal parameters are plotted as a function of the decay rate. Results shown were computed for the CoG model.

Higher decay rates 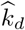 require a higher fraction of mitochondria to be motile in order to maintain optimal health. It should be noted that experimental quantification of neuronal mitochondria indicates that roughly 60 − 90% of mitochondria are in a stationary state [30], in keeping with the optimal values observed for cm-long axons (Fig. 5b,d).

For a fixed fraction of stopped mitochondria, low values of *N*_*s*_ (the number of stops made by a protein during its journey along the axon) correspond to a system with very rare interchange between the stationary and motile populations. Values that are too low result in little opportunity for healthy proteins newly synthesized in the soma to be delivered to the demand sites. By contrast, high values of *N*_*s*_ represent a system with very frequent interchange between the two populations. Overly frequent stopping events increase the probability that a protein in a healthy proximal demand site will be picked up by a retrograde motile mitochondrion and carried back into the soma for recycling. In Supplemental Figure S3, we show that forbidding retrograde fusion events in the Space Station model removes this optimum and allows higher health levels at high values of *N*_*s*_.

Similar behavior with respect to *f*_*s*_ and *N*_*s*_ is seen for the normalized health of the last region (Fig. 5c-d). The optimum stopped mitochondria fraction and number of stopping events are both shifted to slightly lower values in this case, making it more likely that a protein can successfully reach the most distal region without protracted stops along the way.

Figure 5e shows the optimal value for the stationary fraction *f*_*s*_ over a range of dimensionless decay rates 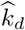. The optimum fraction of mitochondria in the stationary pool is high for very long-lived proteins. In the limit of no decay 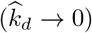, all mitochondria can be kept permanently at the demand sites to maximize the health of the system. In the opposite limit of rapidly decaying proteins, the optimum stationary fraction for the average health approaches *f*_*s*_ ≈ 50%, indicating that mitochondria should be split evenly between motile and stationary pools. For the health of the most distal region, the optimal stationary fraction is slightly lower to allow for more motile mitochondria capable of reaching the end of the domain without extensive decay of their protein content.

In Figure 5f, we see that when the decay rate is relatively low 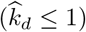, with many proteins surviving a back and forth journey down the axon, optimum mitochondrial maintenance is achieved in the range *N*_*s*_ = 2 − 4 stopping events. This optimal *N*_*s*_ value is not sensitive to specific values of the decay rate, number of demand sites, or total mitochondrial content. It relies only on the assumption of relatively slow protein decay, and the premise that protein exchange is blind to the current health or position of the mitochondria. We note that physiologically relevant values of the dimensionless protein decay rate are expected to fall in the regime 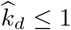. A similar range for the optimal number of stopping events *N*_*s*_ is obtained when considering the health of the most distal demand region only.

Interestingly, the optimum values of *f*_*s*_ and *N*_*s*_ remain largely unchanged in a model with branched tree-like geometries (see Supplemental Material and S2 Fig). Our model thus robustly predicts that the transition of mitochondrial components from motile to stationary organelles at demand sites is expected to be rare (with only 2 − 4 stopping events during the entire journey along the axon), regardless of the specific model, the parameters chosen, or the relative contribution of different sites to the health of the system.

## Effect of Local Translation

While distal protein synthesis plays a role in maintaining and modulating local protein levels in neuronal projections [68], it is unclear to what extent the proteins determining mitochondrial health rely on this mechanism. Approximately 30% of neuronal mRNA transcripts have been identified outside the cell body [41]. For some transcripts that do exhibit local translation, severed axons have been observed to have only 5% total protein synthesis capacity relative to the soma [2]. Although we focus on the long-range delivery of protein components to maintain mitochondrial health, in this section we explore the effect of low levels (5 − 30%) of local protein synthesis that may contribute to mitostasis in neurons.

In our simple model for mitochondrial maintenance, we incorporate a term corresponding to local translation and insertion of proteins specifically into the stationary mitochondria. The total rate of protein incorporation at a demand site is assumed to be proportional to the number of stationary mitochondria at that site, replacing Eq. 2b with

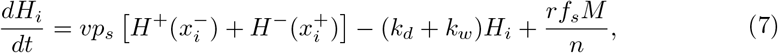

where the stationary mitochondria at steady state are expressed in terms of *f*_*s*_ via Eq. 4.

The total flux of proteins delivered into the axon from the soma varies with the fraction of mitochondria in the motile population. For a given total number of mitochondria *M*, the maximum soma-derived flux is given by *k*_p,max_ = *Mv/*(2*L*). We assume the rate, per mitochondrion, of distal protein manufacture throughout the axon to be equation to a fraction *α* of this somal production, setting *r* = *αk*_p,max_*/M*. The rate at which new proteins are incorporated into mitochondria stationed at each demand site is then given by *rS*.

Figure 6(a,b) shows the effect of a modest level of local translation with *α* = 10%, for high degradation rate 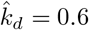. The primary difference compared to the model without local translation (Fig. 5) is the increased health at high values of *f*_*s*_ (large fraction of stopped mitochondria) and at low values of *N*_*s*_ (less frequent exchange between the motile and stationary populations). The optimum value for the fraction of stopped mitochondria is only slightly increased, to *f*_*s*_ ≈ 0.6. Notably, the optimum number of stopping events (*N*_*s*_ ≈ 2 − 4) remains robust to the introduction of low levels of local translation, indicating that infrequent exchanges between stationary and motile populations remain optimal for mitochondrial maintenance.

**Fig 6.**
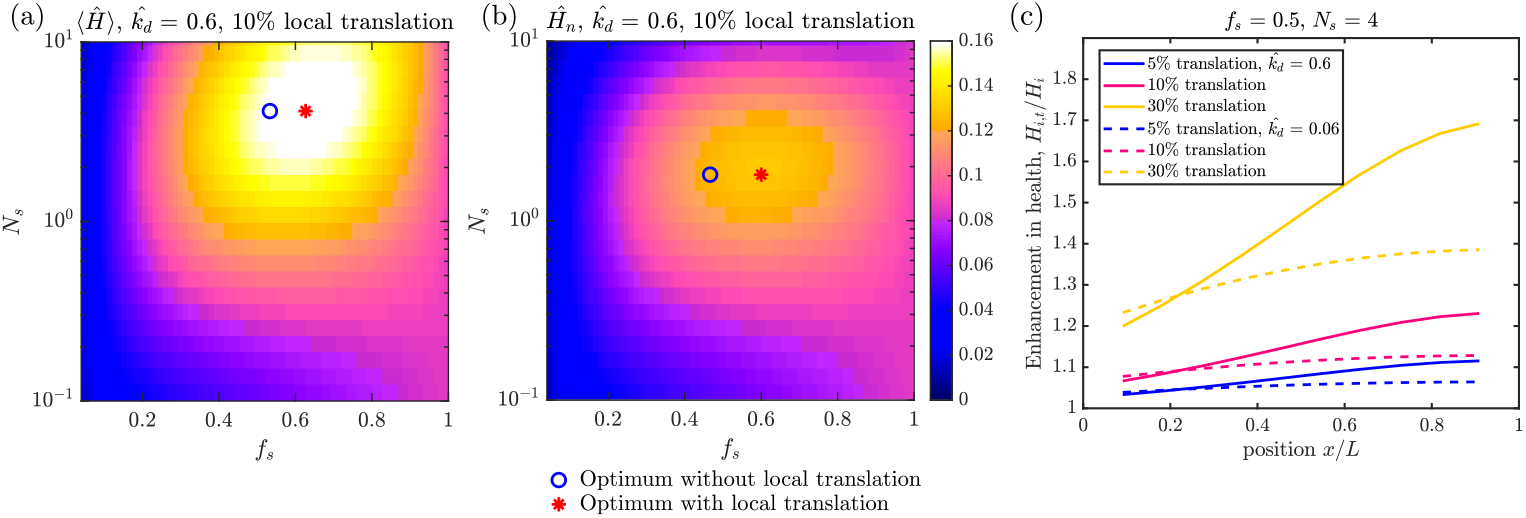
Mitochondrial health in the presence of local translation. (a) Average health across all demand regions as a function of fraction of stopped mitochondria (*f*_*s*_) and number of stopping events (*N*_*s*_), for high decay rate 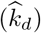 and local translation level *α* = 10%. Markers show the optimal parameter values with (red asterisk) and without (blue circle) local translation. (b) Corresponding mitochondrial health at the most distal demand site. (c) Enhancement of health levels at each demand site in the presence (*H*_*i,t*_) versus absence (*H*_*i*_) of local translation, for one set of transport parameters, (*f*_*s*_ = 0.5, *N*_*s*_ = 0.4), two decay rates 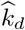, and three different local translation levels (*α*).

In Figure 6(c), we plot the enhancement due to local translation at each of the demand sites (*H*_*i*,t_*/H*_*i*_), for a fixed set of transport parameters close to the optimum value. Unsurprisingly, the effect of local translation is greatest when the dimensionless decay rate is highest, allowing fewer proteins to survive transport to the distal tip of the domain. The biggest effect of local translation is observed on mitochondrial health at the most distal sites. Local translation rates that reach 30% of maximum somal production (per mitochondrion) are sufficient to double the mitochondrial health in the last demand site for high 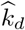 (corresponding to a protein decay time of 4 days and a centimeter-long axon). At these high local translation levels, the optimum in the *f*_*s*_ parameter disappears (S9 Fig), as it becomes advantageous to keep all mitochondria stationed at the demand sites and forgo transport from the soma entirely. However, modest levels of distal translation, similar to those measured for some transcripts in axon severing experiments [2], have a more limited effect on overall health and do little to shift the optimal transport parameters. In order to focus primarily on the interplay between transport and mitostasis, we therefore exclude local translation from our model in all subsequent results.

## Variability of mitochondrial health

Our calculations thus far quantify the mean-field steady-state levels of mitochondrial health in high-demand axonal regions. However, the instantaneous health at each site is inherently variable due to the stochasticity of mitochondrial dynamics. This variability arises from fluctuations in both the number of mitochondria and the health per mitochondrion in each region. A robust cellular distribution system may require high levels of mitochondrial health to be maintained over long time periods, rather than tolerating highly fluctuating health levels. At the same time, variability in the distribution of mitochondria may allow for a more rapid response to changing energy demand at the expense of decreased precision in mitochondrial positioning [35]. In neuronal dendrites, mitochondrial content in localized regions has been shown to fluctuate around a well-defined steady-state level [23], indicating that robust maintenance of healthy mitochondria in those regions (low fluctuations compared to average quantities) may be advantageous to the cell.

To understand the effect of stochasticity in our model, we turn to agent-based simulations. These simulations give the same average steady-state mitochondrial health as the mean-field calculations discussed previously (Fig. 2). We define the total health variability of our system (*σ*_*H*_) as the standard deviation in mitochondrial health per region ⟨*H*⟩, calculated over many iterations of the system after it has reached steady state. Although they exhibit the same average health levels for comparable parameter values, the normalized variability 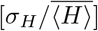 is very different for the two maintenance models (Fig. 7). Specifically, the CoG model has an additional source of variance due to fluctuating numbers of mitochondria per region, whereas the space-station model has a fixed number of mitochondria at each demand site. In the CoG model, replenishment of protein content occurs in whole-mitochondrion chunks, with transient periods of missing mitochondria in the region until a new one arrives to take its place. This discreteness in protein turnover becomes more pronounced when there are fewer mitochondria per region, as seen in Fig. 7. For the SS model, variability arises from how often motile mitochondria exchange their protein contents with stationary ones, and becomes particularly high when the number of motile mitochondria is small (high *f*_*s*_). Because the protein levels are a continuous quantity, the SS model allows for more extensive equilibration across the many mitochondria in the system, and thus exhibits lower fluctuations than the CoG model. This difference between the models becomes even more pronounced at lower decay rates (S6a Fig), when most mitochondria in the SS system have similarly high protein levels but the fluctuations in mitochondria number for the CoG model are still present. It should be noted that while the SS model in our simulations fills up demand sites with stationary mitochondria uniformly (same *S* for all sites), we have also considered an alternate approach where the permanently stationed mitochondria are initially distributed onto demand sites at random. This random initial distribution does not significantly affect the variability in mitochondrial health at the demand sites (S6b Fig).

**Fig 7.**
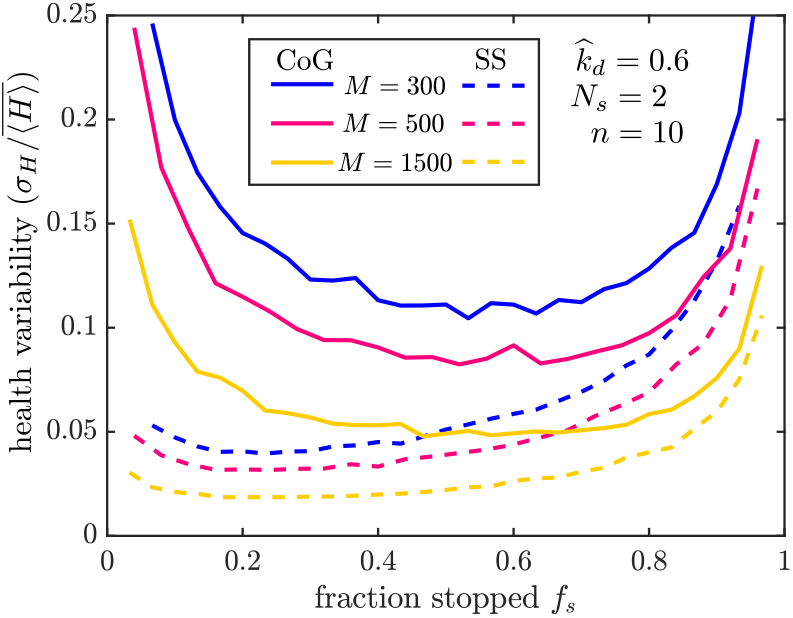
Variability of mitochondrial health in different maintenance models. Plotted is the standard deviation in health per region (*σ*_*H*_) divided by its average value 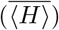, across 1000 iterations of stochastic simulations. Results are shown for 300 (blue), 500 (magenta), and 1500 (yellow) average mitochondria in the domain. Solid lines correspond to simulations of the CoG model and dashed lines to the SS model. All simulations used parameters 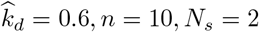. Corresponding plots are provided in Supplementary Material to show the effect of lower decay rates (S6a Fig) and of random stationary mitochondria distribution in the SS model (S6b Fig).

Overall, the SS model maintains more stable health levels in individual demand sites by allowing for equilibration in protein content rather than removal of discrete mitochondria. This maintenance system is thus expected to be more robust during periods of consistent metabolic demand in specific regions. By contrast, the CoG maintenance mechanism gives rise to higher variability that could allow for more rapid redistribution of healthy mitochondria in systems with time-varying demand site positions.

The difference in variability between the two models is a metric which could serve to distinguish the contribution of each to mitostasis in neurons. Experimental data in embryonic fibroblasts indicates that knock-down of the genes responsible for mitochondrial fusion increases heterogeneity in the mitochondria protein age [65]. Similar experiments in neurons would help establish the role of occasional fusion events in maintaining mitochondrial health.

## Effect of Mitophagy

In addition to utilizing motor-driven transport and transient fusion events to maintain a healthy mitochondrial population, neuronal cells make use of mitophagy – a quality control pathway for selectively recycling damaged or depleted mitochondria. Mitochondria with low membrane potential are recognized by the PINK1/Parkin signaling pathway, which triggers ubiquitination and arrest of mitochondrial transport [30, 40, 69]. The marked mitochondria are then targeted for encapsulation by autophagosomes, which transport them back to the soma for recycling while fusing with proximally located fully acidified lysosomes en route [50, 60]. In essence, this pathway allows for the selective removal of damaged mitochondria from the population and directed transport back to the cell body for rapid recycling of the mitochondrial building blocks.

We introduce mitophagy into our model by allowing mitochondria to enter an “engulfed” state whenever their health drops below a threshold level (*ϕ*). This simplified approach neglects any delays in autophagosome arrival or recognition of the damaged mitochondria, assuming that all engulfment events occur immediately when the threshold is reached. Although a variety of other quality control mechanisms, such as asymmetric fission [70] and health-dependent halting of mitochondrial transport [71], may contribute to mitochondrial homeostasis, we focus here on a single simplified sensing process representing mitophagy.

While in the engulfed state, the mitochondria move exclusively in the retrograde direction [62, 63], without pauses or reversals, and are unable to stop or engage in fusion at the demand sites. For the Space Station model, if a permanently stationary mitochondrion becomes engulfed and departs for the soma, it leaves behind a gap at the demand site that is filled by stopping the next motile mitochondrion passing that site. A more complex response involving a stopping probability upon passing each open site would correspond to an interpolation between the SS and the CoG models. Because mitophagy is triggered by low health in an individual discrete mitochondrion, our mitophagy model cannot be easily described by mean-field analytical calculations. Instead, we employ stochastic simulations to explore the effect of mitophagy on steady-state mitochondrial maintenance. Simulations are carried out for a domain with *M* = 300 mitochondria, *n* = 10 regions, and *f*_*s*_ ≈ 0.53, giving a average of 15 mitochondria per site. This number is within the range that has been observed in paranodal regions [29].

### Limiting mitochondrial production rates

In its simplest form, the mitophagy model increases mitochondrial turnover without changing any other parameters. By providing an additional pathway for mitochondria to return to the soma, without altering the production rate *k*_*p*_, mitophagy will decrease the total number of mitochondria in the domain (Fig. 8a). This model represents the limiting case where the number of mitochondria in an axon is limited by the rate at which they can be produced from the somal population and injected into the proximal axon, regardless of how much total mitochondrial material is available. We consider the effect of mitophagy on the overall health of axonal mitochondria in this regime, comparing both the SS and CoG model. We start with the case of no mitophagy (*ϕ* = 0), considering a vertical slice of the plots in Fig. 5 corresponding to a fraction of mitochondria in the stationary state: *f*_*s*_(*ϕ* = 0) = 0.53. We proceed to increase the protein threshold *ϕ*, corresponding to more frequent mitophagy events, while holding constant all other parameters (including production rate *k*_*p*_, number of stops *N*_*s*_, restarting rate *k*_*w*_ in the CoG model, and maximum number of mitochondria that can be stationed at each site, *S*, for the SS model). The effect of increasing mitophagy on mitochondrial health can be considered by defining the health in each demand site, normalized by the total mitochondrial content per site in the absence of mitophagy (*M* ^(0)^). Specifically, we define the health of the *i*^th^ demand site as

**Fig 8.**
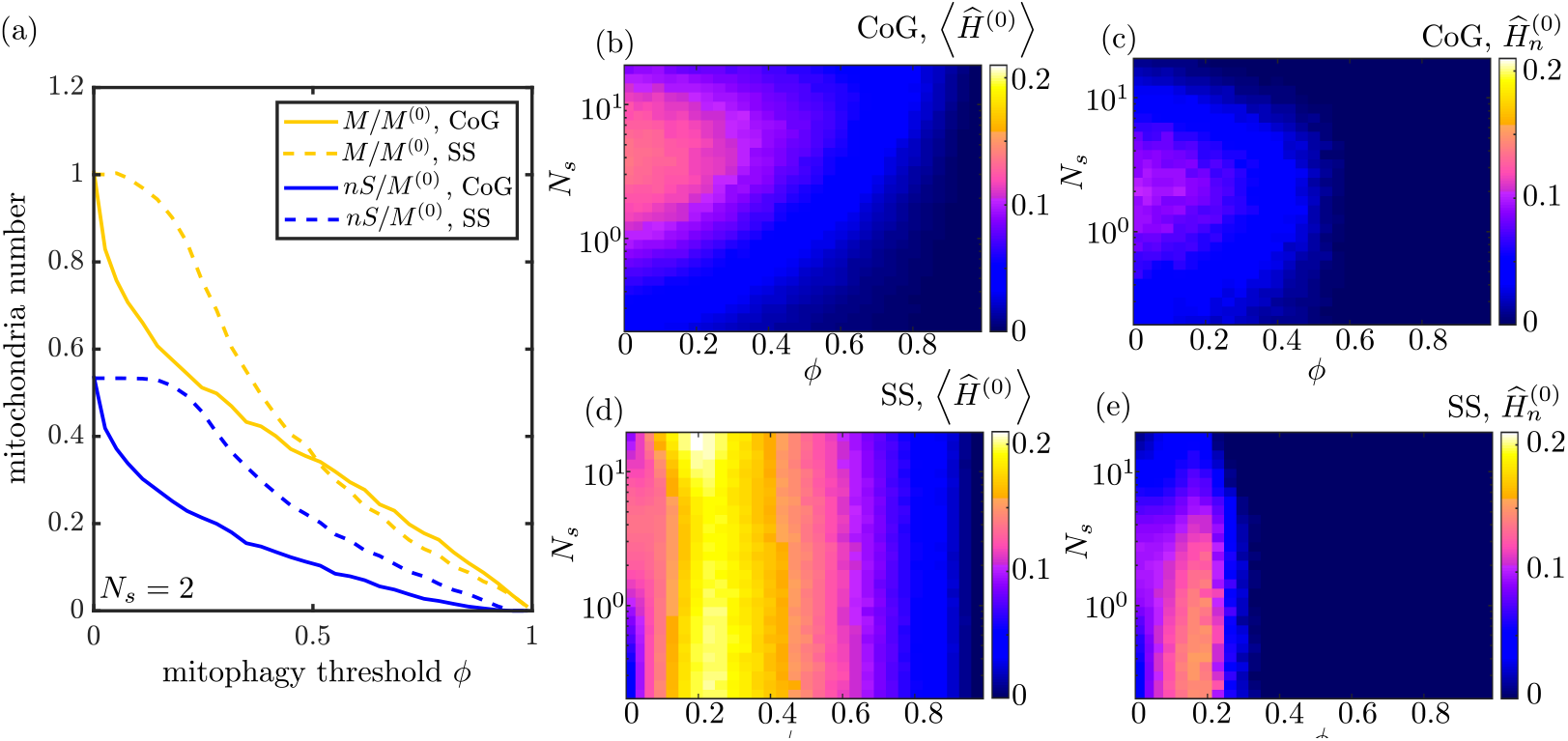
Effect of mitophagy, at fixed production rate *k*_*p*_. (a) The total number of mitochondria in the domain (yellow) and the number of stationary mitochondria (blue) are plotted as a function of the mitophagy threshold, for both the CoG model (solid lines) and the SS model (dashed lines). Mitochondria quantities are normalized by the steady-state number of mitochondria in the absence of mitophagy. (b) Mitochondrial health for the CoG model, averaged over all demand sites, plotted as a function of mitophagy threshold and number of stopping events *N*_*s*_. (c) Health at most distal demand site, for the CoG model. (d-e) Analogous plots of average health and distal site health for the SS model. All plots assume mitochondrial production rate does not change with increased mitophagy, and correspond to fraction of mitochondria stopped *f*_*s*_(*ϕ* = 0) = 0.53 in the absence of mitophagy.

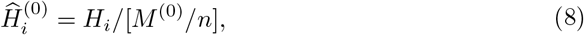

and the average health across all sites as ⟨ *Ĥ* ^(0)^ ⟩. Unlike our previous normalization (Eq. 1), this quantity no longer corresponds to the health per mitochondrion, because the advent of mitophagy would decrease the total number of mitochondria in the domain. Instead, we use this health metric to understand how the total amount of mitochondrial protein changes at each demand site when new mitochondrial production is not upregulated to keep up with the enhanced turnover due to mitophagy. The alternate case, where total mitochondrial content is kept constant, is considered in the subsequent section.

A seen in Fig. 8b,c, the introduction of mitophagy in the CoG model decreases the mitochondrial health at the demand sites. This is a result of a decrease in the average number of mitochondria stationed at each site (Fig. 8a), as mitophagy causes more mitochondria to leave the domain without increasing the production rate *k*_*p*_. By contrast, in the SS model modest levels of mitophagy will actually increase the average mitochondrial health at the demand sites (Fig. 8d). For this maintenance system, the number of stationary mitochondria does not change significantly for *ϕ* ≲ 0.2 (see Fig. 8a). However, these low levels of mitophagy remove some of the most unhealthy motile mitochondria from the population capable of fusing, thereby increasing the average health of all the remaining mitochondria in the domain. Notably, a mitophagy threshold of *ϕ* ≈ 0.2 optimizes both the average health of stationary mitochondria and the health of the most distal site in the SS model (Fig. 8d-e).

The difference between the CoG and SS model in the presence of mitophagy reflects, in part, the different variability of stationary mitochondria. In the SS model, the mitochondria are more homogeneous in terms of their protein content (Fig. 7). Because mitophagy removes individual organelles whose health drops below a particular threshold, the homogenization arising from fusion events makes it less likely that a stationary mitochondrion will drop below threshold and become engulfed. As a result, low levels of mitophagy have a much smaller effect on both the number of stationary mitochondria and the total number of mitochondria (engulfed, stationary, and motile) in the SS model. For this model, only those mitochondria moving retrograde and already near the somal region become engulfed at low *ϕ* values. These depleted mitochondria are prevented from fusing with stationary mitochondria in the proximal demand regions, thereby increasing the overall health of the system without substantial change to total mitochondrial number.

### Limiting mitochondrial content

One of the presumed benefits of the mitophagy pathway is its ability to more rapidly return damaged mitochondria to the soma, permitting their components to be recycled for future use [29]. We therefore consider again the regime where the system is limited not directly by the production rate, but by the total amount of available mitochondrial components. In this case, mitophagy causes an increase in the production rate *k*_*p*_, as the mitochondria on average return to the soma more quickly for recycling. We implement this model by rescaling all protein levels by the total number of mitochondria (including engulfed, stationary, and motile) in the domain. Health levels are normalized by the actual number of total mitochondria at steady state [*Ĥ*_*i*_ = *H*_*i*_*/*(*M/n*)], thereby removing the effect of decreased mitochondrial population in the presence of autophagy. This approach is equivalent to explicitly fixing the total amount of mitochondrial content available, as demonstrated in Supplementary Material (S7 Fig).

In this regime, both the SS and CoG maintenance models allow an increase in mitochondrial health with intermediate levels of mitophagy (Fig. 9b-e). The normalized health of the most distal region (*Ĥ*_*n*_) is increased by about 90% for the SS model and 40% for the CoG model, when compared to the case of zero mitophagy (evaluated at the optimum value of *N*_*s*_ for each *ϕ* value). As before, the optimal mitophagy threshold is in the range *ϕ* ≈ 0.1 − 0.2 for both models. The average health of stationary mitochondria improves by 80% for the SS model and 20% for the CoG model at these optimal mitophagy thresholds. While higher mitophagy levels can improve average regional health still further, they do so at the expense of a severe drop in the health of the most distal region, as very few mitochondria are able to reach that region when mitophagy occurs too early in the domain.

**Fig 9.**
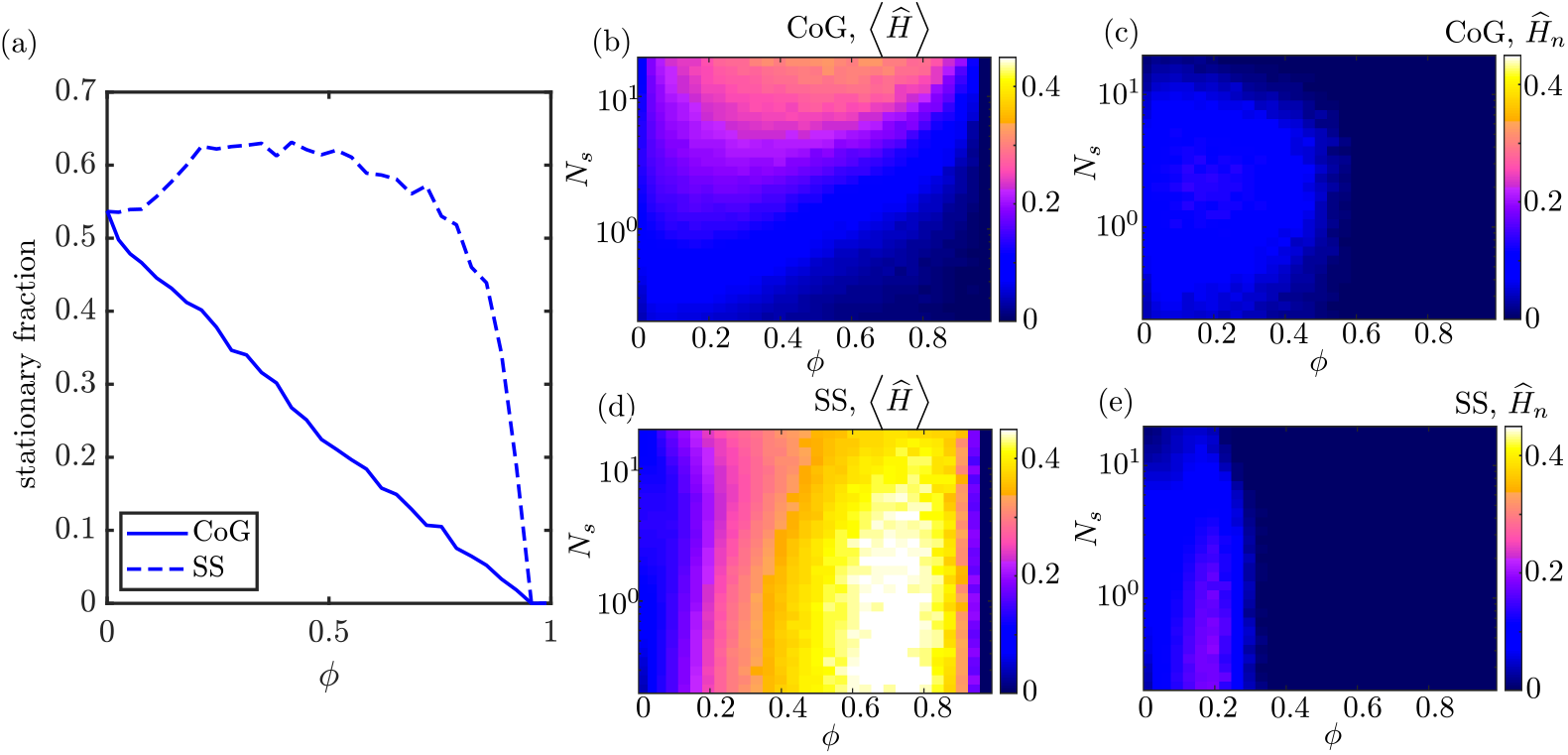
Effect of mitophagy when total mitochondrial number is limited. (a) Fraction of mitochondria in the stationary state as a function of increasing mitophagy threshold, for the CoG model (solid line) and the SS model (dashed line), with parameters set such that *f*_*s*_ = 0.53 at *ϕ* = 0. (b) Mitochondrial health, normalized by the total number of mitochondria per site, averaged over all demand sites, for the CoG model. (c) Normalized mitochondrial health at most distal demand site, for the CoG model. (d-e) Analogous plots of normalized average health and distal site health for the SS model. Normalizing by total mitochondrial number is equivalent to a system where the total mitochondrial content is held fixed with the onset of mitophagy (S7 Fig).

Even when normalizing by total mitochondrial content, the SS model has a significant advantage over the CoG model. Namely, because the restarting rate *k*_*w*_ is fixed for each row in the plots of Fig. 9, the fraction of mitochondria in the stationary state drops substantially in the CoG model as mitophagy is introduced (Fig. 9a). This reflects the fact that stationary mitochondria are preferentially engulfed in this model, so that even if we normalize by the total mitochondrial content, a smaller fraction of mitochondria remain at the demand sites. In the SS model, by contrast, equalization of mitochondrial health by transient fusion events implies that stationary mitochondria are no more likely than motile ones to become engulfed, and the fraction of mitochondria in the stationary state remains constant or even increases slightly for a broad range of *ϕ* values.

It is reasonable to suppose that a cell employing the CoG strategy might have compensatory mechanisms to adjust the fraction of the mitochondrial pool that is stationed at demand sites in the presence of mitophagy (perhaps by increasing stopping probability *p*_*s*_ or decreasing the restarting rate *k*_*w*_). In order to assess the relative advantage of the two models, we therefore consider the maximal mitochondrial health each of them can achieve at a given mitophagy level, if allowed to freely adjust the other model parameters. We assume the total mitochondrial pool remains fixed and therefore again normalize the results by *M/n* as in Fig. 9.

As the mitophagy threshold is raised (with freely adjustable transport parameters), the increase in average health over all the demand regions can be substantial — up to 4-fold for the CoG model and 5-fold for the SS model (Fig. 10a). However, the parameters that give such high average health result in most of the protein content being concentrated in the first few demand sites, with the health of the most distal site dropping to zero. This regime corresponds to rapid turnover of motile mitochondria that undergo mitophagy before they ever reach the distal regions. Notably, experimental observations have indicated that mitochondrial densities are indeed somewhat lower at distal sites [72, 73]. Given the necessity of supplying the metabolic needs of all individual demand sites, we next consider the parameters necessary to maximize mitochondrial health in the most distal site, rather than averaging over the entire domain.

**Fig 10.**
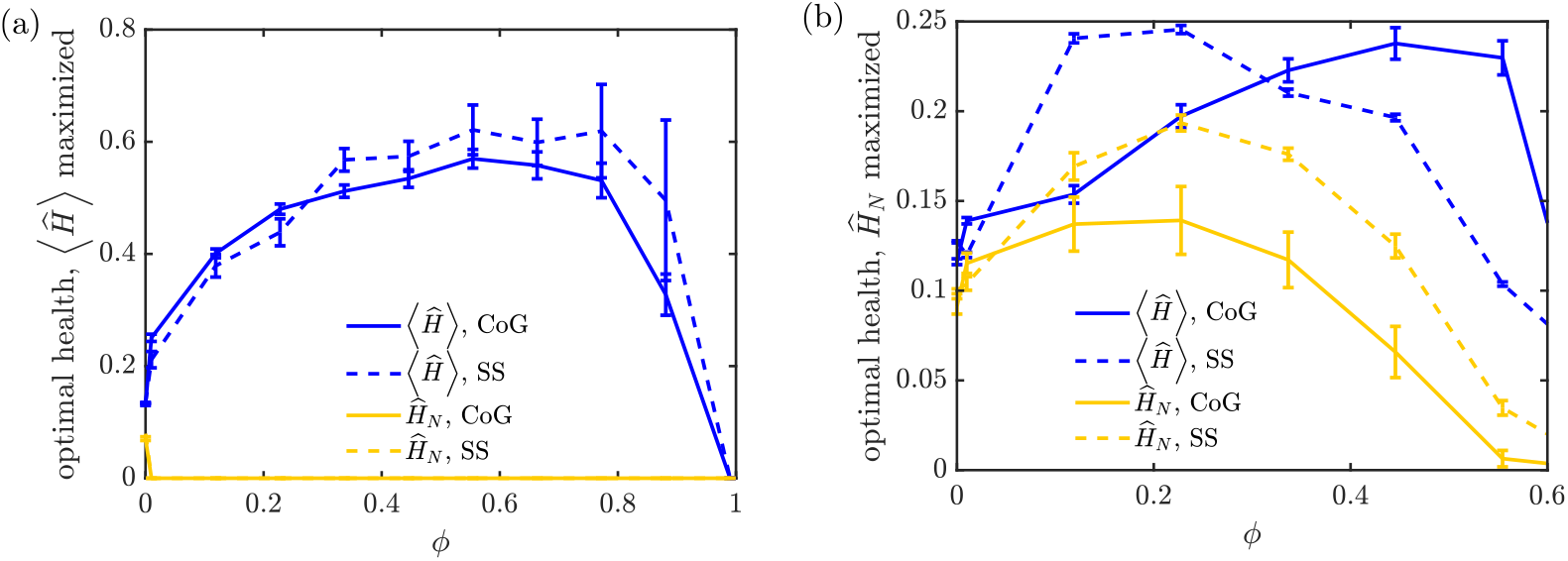
Optimal performance of mitochondrial maintenance models in presence of mitophagy. (a) For each mitophagy threshold *ϕ*, the parameters *f*_*s*_, *N*_*s*_ are optimized to give the maximum normalized average health: ⟨*H*⟩. The resulting normalized average health (blue) and normalized last region health (yellow) are plotted for the CoG model (solid) and SS model (dashed). (b) Analogous plots, with parameters adjusted to maximize the normalized last region health *Ĥ*_*n*_ for each mitophagy threshold. All health levels are normalized by total mitochondrial content per region (*M/n*), corresponding to a system where the total amount of mitochondrial material is limited. Error bars show standard error of the mean from 10 replicates. Fixed parameters are 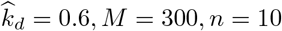. A corresponding plot of optimized health for low decay rate 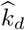 is provided in Supporting Information Fig. S8

If we optimize over the health of the most distal site (Fig. 10b), we see as before that this distal health increases by about 90% in the SS model and about 40% in the CoG model, compared to the case of no mitophagy. These values represent the maximal distal site health that can be achieved by each model if the parameters *f*_*s*_ and *N*_*s*_ are allowed to vary freely, within the constraint of constant *ϕ*, 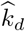, *M, n*. At the optimal mitophagy threshold (*ϕ* ≈ 0.2), the average regional health is also increased, in both models, by up to a factor of two. However, the CoG model allows for greater average health at still-higher mitophagy levels, indicative of its tendency to distribute mitochondria unevenly throughout the domain when mitophagy is present, so that proximal sites gain many healthy mitochondria while distal health are left under-supplied.

We conclude that introducing mitophagy to selectively recycle unhealthy mitochondria can substantially improve mitochondrial health throughout high-demand regions of the domain. This improvement is maximized by allowing mitochondria to become engulfed when their health level drops below approximately 20% of its initial value. It should be noted that this optimal value of *ϕ* is specific to the decay rate 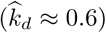 used in these simulations. This dimensionless decay rate corresponds to a domain length of about 10 cm and protein lifetime of about 4 days, so that there is a substantial amount of protein decay by the time a mitochondrion traverses the entire domain. Lower values of 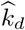 move the optimal *ϕ* to higher values, rising to an optimum value of *ϕ* = 0.6 for 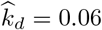, corresponding to a shorter 1 cm axon (S8 Fig). This shift reflects the fact that less degradation occurs during the journey of a mitochondrion along the axon, making it feasible to trigger mitophagy at higher health thresholds while still allowing mitochondria to reach the last demand site. An interesting consequence of this model is that longer neuronal projections require mitophagy to be delayed to more extensive decay levels in order to permit at least partially functional mitochondria to reach the most distant demand sites.

## Discussion

The models described above constitute a quantitative framework for mitochondrial maintenance in extended cellular regions such as neuronal axons. Neurons face unique challenges in mitochondrial homeostasis, both because of the need to transport material from the cell body through long cellular projections and because of their spatially heterogeneous metabolic needs. Balancing these constraints requires positioning stationary mitochondria in specific regions of high metabolic demand, while also retaining a population of motile mitochondria to transport the components needed for mitochondrial health. Our work differs from prior studies of mitochondrial maintenance [51, 52] by taking into account these spatial constraints unique to neurons. At the same time, we diverge from prior models of transport and spatial distribution for neuronal components [35] by incorporating continuous degradation and focusing on the steady-state health at specific localized sites.

Our results highlight similarities and differences between two main models of mitostasis: the ‘Changing of the Guard’ and the ‘Space Shuttle’ model. In the absence of selective recycling processes, the two models are shown to be equivalent in their average steady-state behavior. Analysis of the mean-field model for both mechanisms shows that in extended projections, with many demand sites and large numbers of mitochondria, the efficiency of the maintenance mechanism depends primarily on three dimensionless parameters. These parameters are (1) 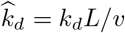, which sets the extent of health decay while a mitochondrion crosses the domain, (2) *f*_*s*_ = *nS/M*, the fraction of mitochondria in the stationary pool, and (3) 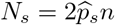, the typical number of stopping events for a mitochondrial component during its round-trip journey through the projection. Optimal mitochondrial health in high-demand sites is achieved at intermediate values of both *f*_*s*_ and *N*_*s*_. The optimal fraction of stationary mitochondria lies in the 50 − 80% range, depending on the precise value of the dimensionless decay rate. This range is compatible with experimental observations which imply that 60 − 90% of axonal mitochondria are stationary [30]. For mature cortical axons, the observed stationary fraction has been observed to be relatively high at 95% [74], possibly hinting at an even slower effective degradation rate of mitochondrial cargo.

Furthermore, our model makes the prediction that, for a broad physiologically relevant range of decay rates, very few stopping events (*N*_*s*_ ≈ 2 − 4) are required for optimal maintenance. This result is robust to the introduction of low levels of local translation (Fig 6) or expansion of the model to a branched tree-like morphology (S1 Fig). The low optimal value of *N*_*s*_ implies that both switching of mitochondria between motile and stationary states and transient fusion events of axonal mitochondria should be very rare – happening only a few times during the entire journey of a mitochondrion down the axon. Due to the inherent difficulty of experimentally tracking motile mitochondria over long time periods, this result is consistent with the fact that stationary mitochondria appear to be nearly permanent and interchange between stationary and motile pools is rarely observed [59, 75, 76].

Differences between the CoG and the SS models become evident when considering the variability of mitochondrial health at demand sites over time and between different cells. Transient fusion and fission events in the SS model allow extensive equilibration of health levels between individual mitochondria, resulting in much lower variance in the health of entire regions. On the one hand, such decreased variance implies a more robust maintenance system, which avoids large fluctuations in metabolic health at a high-demand site as individual mitochondria enter or leave the stationary pool at that site. On the other hand, greater fluctuations in the CoG model may imply an enhanced responsiveness to changing conditions (*eg:* changing spatial distribution of metabolic demand). A close relationship between fluctuation magnitudes and response to external driving is a key feature of physical systems dominated by thermal fluctuations [77].

Analogous fluctuation-response relationships have also been proposed to underly biological systems [78, 79], such as gene networks, where a greater degree of responsiveness to external signals goes hand in hand with higher fluctuations at steady-state [80, 81]. Although detailed exploration of temporal response by mitochondrial maintenance systems lies outside the scope of this paper, the behavior of these models in the presence of time-varying metabolic demand serves as a potentially fruitful area for future work.

A biologically relevant consequence of the different fluctuation magnitudes for the two maintenance models is their differing response to the incorporation of selective mitophagy of damaged mitochondria. In a system where mitochondrial production remains constant, introducing low levels of autophagy causes a substantial drop in the number of stationary mitochondria at each demand site in the CoG model. For the SS model, by contrast, equilibration of mitochondrial health implies that the number of both stationary and total mitochondria remains relatively unchanged at low mitophagy thresholds. As a result, moderate autophagy levels improve the overall mitochondrial health at demand sites in the SS model while monotonically decreasing health in the CoG case.

Because mitophagy allows mitochondria to be recycled more rapidly, we also consider the steady-state health normalized by total number of mitochondria in the domain. This approach is equivalent to adjusting the mitochondrial production rate in such a way that the total number of mitochondria remains fixed even in the presence of autophagy. In this case, an optimal autophagy threshold is observed for both models, although the SS model still allows for the greatest (nearly 2-fold) increase in the mitochondrial health at the most distal demand site. Overall, the extensive mixing of mitochondrial contents permitted by transient fusion and fission events allows for improved health of localized mitochondria in the presence of recycling via selective mitophagy. The optimal mitophagy threshold depends on the domain length and protein decay rate, with an optimal value of *ϕ* ≈ 0.2 for both models in the case of 10 cm axons and 4-day mitochondrial protein decay times.

The models described in this work are, of necessity, highly simplified. The model design aimed to highlight key challenges of mitochondrial maintenance in long neuronal projections and some of the fundamental strategies that can be utilized by the cell to meet those challenges. In reality, neuronal mitostasis may include a variety of complicating factors, many of which remain poorly understood. For example, our model incorporates the simplest possible sensing and response mechanism for mitochondrial health — dropping below a critical health level prevents further exchange events and forces a return to the soma for recycling. Other contributing mechanisms may include asymmetric fission that concentrates health factors in one of the resulting organelles [52, 70], local recycling [60] and formation of new mitochondria, or a sensing mechanism that regulates the ability of mitochondria to stop as a function of total ATP production by other mitochondria in the region [31, 82]. Furthermore, a sensing mechanism that would prohibit fusion of retrograde mitochondria while allowing frequent fusion of anterograde ones would also increase overall health levels in the domain, though having little effect on the health of the most distal site (S3 Fig). Although such mechanisms would allow for more efficient mitochondrial maintenance, we focus here on the simplified case of purely somal mitochondria production and no additional sensing that would differentiate the behavior of stationary or motile organelles. Additionally, we employ the simplifying assumption of a constant decay rate *k*_*d*_ for both stationary and motile mitochondria. More complicated models where the decay in mitochondrial health depends on local metabolic activity could be incorporated within the framework of this model in future work.

In addition to the two transport and exchange mechanisms described here, local protein translation may also contribute to mitochondrial maintenance in distal regions [3–5]. However, the extent to which mitochondria are able to produce their full complement of proteins by local synthesis is largely unknown. General measurements of local translation indicate that only about a third of transcripts are found outside the cell soma, and that for individual transcripts the protein synthesis capacity of the axon may be only a small fraction of somal production [2, 41]. We incorporate a small contribution from local translation in our model, and find that the primary outcomes – the optimal fraction of stopped mitochondria and the small optimal number of stopping events – remain unchanged (Fig. 6). More extensive local synthesis could drive the system to place all mitochondria in the stationary population (S9 Fig). However, it should be noted that even a small set of mitochondrial proteins that rely on somal translation would necessitate the transport and exchange mechanisms described here. To focus on the interplay between transport and maintenance, we primarily explore system dynamics in the absence of local translation. Our assumption that key components of mitochondrial health require long-range transport from the soma is partly substantiated by evidence that mitochondrial aging increases with distance from the soma [65], implying an important role for long-range transport.

For the most part, we consider the simplest possible system geometry, with a single linear projection rather than more complicated branched axonal structures. We find that our results with this simplification can also be applied to symmetric branched geometries, further underscoring the importance of the effective transport and exchange parameters defined here. Our models also assume the metabolic demand in a neuron is concentrated at discrete high-demand regions. While this is an over-simplification of the complex distribution of neuronal metabolism, it serves as a limiting case for spatial heterogeneity that requires the formation of both stationary and motile pools of mitochondria.

Studies in non-neural globular animal cells show that mitochondrial maintenance relies largely on fusion and fission to maintain the health of the overall population of mitochondria [42]. In neuronal axons, fusion events are thought to be relatively rare, and it seems likely that mitostasis actually involves a hybrid of the ‘Changing-of-the-Guard’ and the ‘Space-Shuttle’ mechanisms. Our separate exploration of the two models highlights the similarity of both in maintaining the average mitochondrial health. However, we have also shown that the differing fluctuations dictated by the two models can alter their response to rising levels of mitophagy, implying that the introduction of occasional localized fission and fusion events can prove beneficial. Experimental studies that explore the effect of knocking down fusion proteins on the overall age of mitochondrial proteins, such as have already been carried out for non-neuronal cell types [65], would help disentangle the contribution of the SS and CoG mechanisms to mitostasis in neurons.

The interplay of transport and localization is a general principle for maintaining homeostasis under the constraints of highly extended geometries and spatially heterogeneous demands that characterize neuronal cells. Our analysis outlines the key parameters that can be tuned to optimize steady-state distributions, with a specific focus on maintaining mitochondrial health in neurons. A key consequence of the models is that optimal mitochondrial health can be maintained with a large fraction of stationary mitochondria and with very rare interchange between the stationary and motile population. This prediction may account for the difficulty of catching such exchanges in live cell imaging experiments, and the general tendency of stationary mitochondria to remain stationary over long periods of observation [74–76]. With the advent of new experimental techniques that allow for long-term tracking of individual mitochondria [59], it would be valuable to quantify the frequency of these rare transitions. Our results also predict a benefit to mitochondrial exchange via transient fusion events, motivating future experimental determination of whether such fusions contribute to maintenance of localized axonal mitochondria. Given the importance of mitochondrial maintenance for neuronal health, developing a quantitative framework that delineates the factors governing mitostasis serves as a critical step towards better understanding of the role of metabolism in neurodegenerative diseases.

## Methods

Source code for all simulations and analytic calculations is available at: https://github.com/lenafabr/mitofusion.

### Table of Parameters

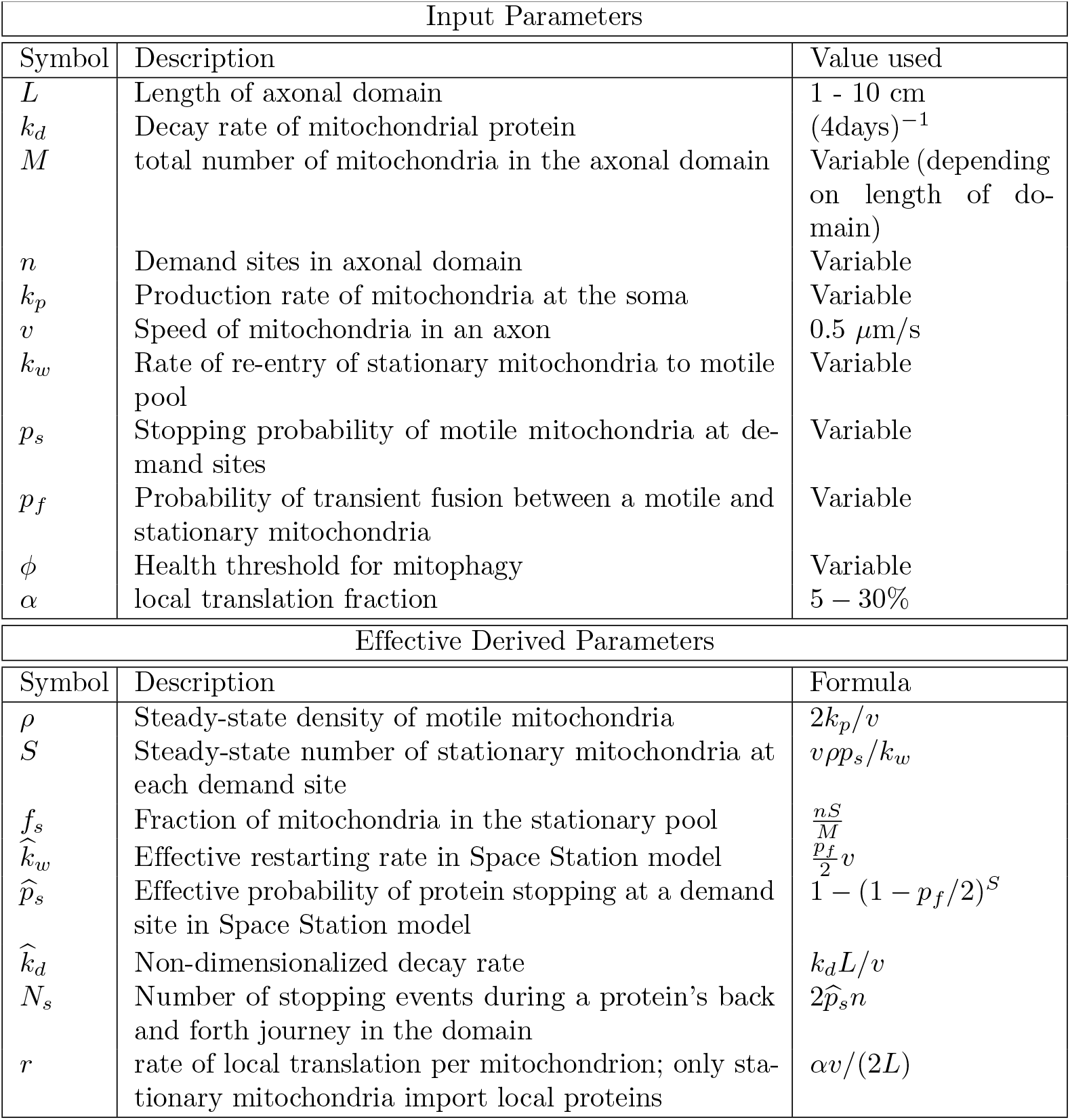

### Steady-state mean-field solution for CoG model

We find the steady-state solution for the mean-field version of the Changing-of-the-Guard model, which treats mitochondrial density and mitochondrial health as continuous fields on a linear segment from *x* = 0 to *x* = *L*. We define the density of anterograde and retrograde motile mitochondria as *ρ*^*±*^(*x*), while the number of stationary mitochondria in each point-like high-demand region *i* is given by *S*_*i*_. These distributions obey the following set of dynamic equations:

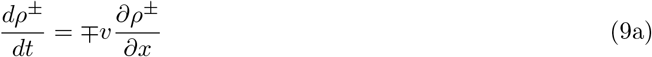

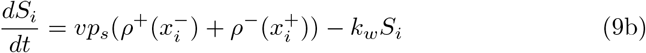

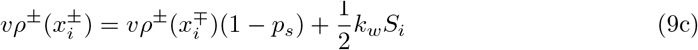

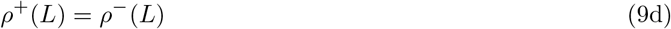

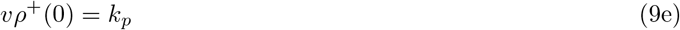

Equation 9a describes the evolution of the mitochondrial density in each interval between consecutive demand sites. Equation 9b governs the number of stopped mitochondria at each discrete demand site, with the first term corresponding to the incoming flux of mitochondria stopping at that site, and the second term corresponding to the restarting of stationary mitochondria. The remaining equations provide the boundary conditions for each interval of the domain between demand sites. Equation 9c enforces conservation of organelles, so that the flux of anterograde mitochondria leaving the site is equal to the combination of those organelles which pass by without stopping and those which restart from a stopped state. Eq. 9d - 9e set the boundary conditions at the distal end (anterograde mitochondria turn around to move in the retrograde direction) and the proximal end of the domain (mitochondrial production).

At steady state, Eq. 9a implies that the linear density of mitochondria is constant within each region between demand sites. Setting Eq. 9b to zero yields the steady-state relation,

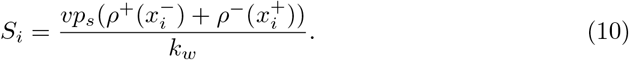

Plugging in to the boundary conditions Eq. 9c-9e then implies that the motile mitochondrial densities are equal and constant everywhere in the domain: *ρ*^+^ = *ρ*^*−*^. We define *ρ* = *ρ*^+^ + *ρ*^*−*^ as the constant steady-state density of all motile mitochondria. From the boundary condition at the proximal end, we can relate this density (as well as the stopped mitochondria at each site) to the rate of mitochondrial production.

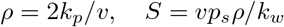

The linear density of mitochondrial health (ie: the sum of health levels for all mitochondria that happen to be present at a given position in the domain) can be described by very similar equations, with additional terms for decay at rate *k*_*d*_. Equation 2 describes the evolution of this field over time.

Setting Eq. 2b-2c to zero at steady-state yields the relations

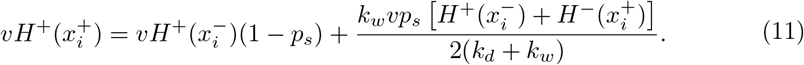

Furthermore, Eq. 2a implies that the steady-state distribution of motile mitochondrial health exhibits an exponential decay across each interval between consecutive demand sites *i* − 1 and *i*. This gives the set of conditions:

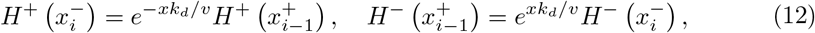

Together, Eq. 11, 12, and 2c - 2e constitute a set of linear equations for the health levels in stationary mitochondria at the demand sites [*H*_*i*_] and the motile health densities on either edge of the demand sites 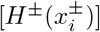. This system of equations is then solved using standard matrix methods.

### Steady-state mean field solution for SS model

A key feature of the Space Shuttle model is that a fixed number of permanently immobile mitochondria (*S*) are stationed at each discrete demand site. These mitochondria are assumed to have a well defined order, but to be positioned arbitrarily close together within the point-like site. We define *H*_*i,j*_ (with 1 ≤*i* ≤*n* and 1 ≤ *j* ≤ *S*) as the health of the *j*^th^ stationary mitochondrion at the *i*^th^ demand site. An anterograde moving mitochondrion encounters each of the stationary ones in order as it passes the demand site. The quantity 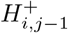 describes the linear health density of all anterograde mitochondria just after they pass the (*j* − 1)^th^ stationary organelle at the *i*^th^ site, and are approaching the *j*^th^ stationary organelle. Similarly, retrograde mitochondria encounter the stationary organelles in reverse order and their health density is given by 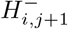 as they approach the *j*^th^ stationary organelle from the distal side. Each time a moving mitochondrion passes a stationary one, there is a probability *p*_*f*_ that they will fuse. When such a fusion event occurs, the health levels of the two mitochondria are averaged together and both are left with a health equal to that average. It should be noted that *H*^*±*^ and 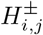 are densities and have units of health level per unit length, while the quantities *H*_*i,j*_ have units of health level.

Motile anterograde mitochondria are produced at the proximal end with rate *k*_*p*_. They then move forward and back throughout the domain, never stopping and instantaneously reversing their direction from anterograde to retrograde once they reach the distal end. The density of motile mitochondria is not affected by the fusion behavior, and evolves according to Equations 9a, 9d, 9e. At steady state, this density must be a constant value *ρ* = (*M* − *nS*)*/L*, equally split between anterograde and retrograde organelles.

The set of mean-field equations describing the mitochondrial health in the SS model is, for 1 ≤ *j* ≤ *S*,

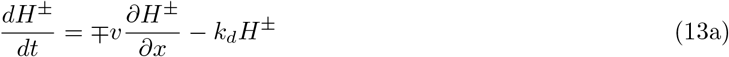

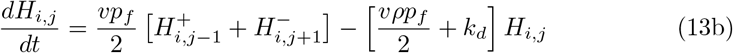

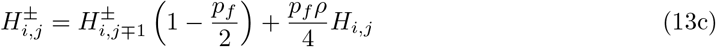

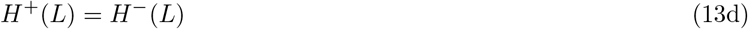

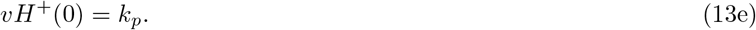

Equation 13b describes the time evolution of the health of each stationary mitochondrion. The first term corresponds to the flux of incoming mitochondrial health multiplied by the probability of fusion (*p*_*f*_) and the probability (1*/*2) that a given protein will stay with the stationary mitochondria after the fusion-and-fission cycle is complete. The second term gives the rate at which health markers leave the stationary mitochondrion through being transferred to the motile partner during a fission and fusion cycle, and through decay. The edge values at each site are defined by 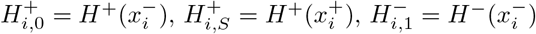, and 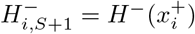, where 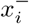 refers to the limit approaching site *i* from the proximal side and *x*^+^ is the limit approaching from the distal side. Equation 13c sets the flux of motile health leaving a stationary mitochondrion equal to the flux of health approaching that mitochondrion from the opposite side and going straight through without transfer, plus the flux of health carried out of the stationary mitochondria as a result of fusion events. Overall, Eq. 13 provides a closed set of equations that can be solved at steady state.

In the case where there is one stationary mitochondrion at each demand site (*S* = 1), the equations above become equivalent the the dynamic equations for the CoG model (Eq. 2), with the restarting rate *k*_*w*_ replaced by an effective rate at which proteins in a stationary mitochondrion leave the organelle via fusion and fission 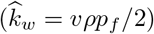. The stopping rate *p*_*s*_ is replaced by an effective rate of a protein ending up in a stationary mitochondrion each time there is a passage event 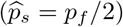. In the case where there are multiple mitochondria per site, the total stationary health at each site is given by 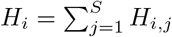. Solving together Equations 13b-13c then yields

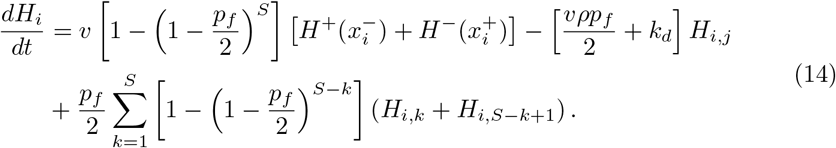

For small values of *p*_*f*_, the last term can be neglected, and the SS model becomes equivalent to the CoG model (Eq. 2b), with the effective stopping rate given by 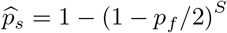, and the effective restarting rate given by Eq. 5. Intuitively, the effective stopping rate is simply obtained by finding one minus the probability that a protein is not left behind in any of the *S* independent fusion events, each of which occurs with probability *p*_*f*_ while passing the multiple mitochondria stationed at a site.

### Discrete stochastic simulations

Discrete stochastic simulations for both the CoG and the SS model were carried out by tracking the positions and health levels of individual point-like mitochondria on a finite linear interval representing a neuronal axon. All length units were normalized by the domain length and all time units by the mitochondrial velocity, such that *L* = 1 and *v* = 1 in the simulations. High-demand sites where mitochondria could stop (in the CoG model) or fuse (in the SS model) were represented as point sites equispaced in the domain. Simulations were propagated forward for 10^5^ time-steps of duration Δ*t* = 10^*−*3^. Each simulation was carried out as 100 independent iterations and the average results are reported.

At each time-step, a new mitochondrion was generated at position 0 with probability 1 − exp(−*k*_*p*_Δ*t*). For the CoG model, each mitochondrion was labeled as being in the anterograde, retrograde, or stationary state. At each time-step, a motile mitochondrion took a step of ± *v*Δ*t*, depending on whether it was in the anterograde or retrogtrade step. An anterograde mitochondrion that reached the domain end (at *L*) was flipped to a retrograde state. A retrograde mitochondrion that reached 0 was removed from the simulation. A stationary mitochondrion restarted with probability 1 − exp(−*k*_*w*_Δ*t*), with its new directionality equally likely to be set to antegrograde or retrograde. Whenever a motile mitochondrion crossed the demand site, it had probability *p*_*s*_ of switching to a stationary state. Each mitochondrion had a continuous health variable associated with it, set to 1 when the mitochondrion was formed and multiplied by exp(−*k*_*d*_Δ*t*) during each simulation time-step.

The SS model was simulated in a similar manner. A fixed number of permanently stationary mitochondria were evenly distributed among the demand sites at the beginning of the simulation. Each time a motile mitochondrion crossed over the demand site, it attempted to fuse with each stationary mitochondrion at that site, in sequence, with probability *p*_*f*_. Whenever a fusion successfully occurred, the health levels of both fusing mitochondria were set to the average of their two health levels. The motile mitochondrion would then proceed to attempt the next fusion or move further down the domain. Fission and fusion events were assumed to be instantaneous.

Mitophagy in the stochastic simulations was implemented by setting a cutoff level *ϕ* for the health of an individual mitochondrion. Whenever a mitochondrion’s health dropped below that cutoff, it was labeled as “engulfed”. An engulfed mitochondrion moved in the retrograde direction (−*v*Δ*t*) on each time-step and was not subject to fusion, fission, or stopping events. Once it reached the proximal end of the domain (position 0) it was removed from the simulation. The total number of mitochondria in the domain (*M*) included these engulfed particles. However, their health levels were not included in any calculations of health at the demand sites. Simulations with mitophagy (Fig. 8, 9, 10) were carried out with *M* = 300.

For S7 Fig, additional stochastic simulations were carried out for a variant of the CoG model with explicitly fixed total number of mitochondria in the domain. Specifically, *M* mitochondria were initially placed uniformly at random in the domain in a motile state. Each time a retrograde mitochondrion reached *x* = 0, it switched immediately into an anterograde mitochondrion with health level reset to 1. No new mitochondria were produced, and the system was allowed to run for 10^5^ time-steps to reach steady-state. The number of mitochondria therefore stayed fixed throughout, regardless of the presence of mitophagy.

### Optimizing performance in the presence of mitophagy

For each of the plots showing the effect of mitophagy in Fig. 8, 9, the effective stopping rate was set to give a particular value of *N*_*s*_. Simulations were carried out with *n* = 10 demand sites, and *M* = 300 average total mitochondria in the domain.

For the data shown in Fig. 8, 9, the fraction of stopped mitochondria was set to *f*_*s*_(*ϕ* = 0) = 0.53 in the absence of mitophagy. In the SS model, a number *f*_*s*_*M*_0_ of permanently stationary mitochondria were placed evenly distributed among all the demand sites. In the CoG model, the restarting rate *k*_*w*_ was set according to Eq. 4, for each value of *N*_*s*_. The production rate was then set according to Eq. 3, to give the desired number of motile mitochondria in the domain at steady-state. The value of *k*_*p*_ was not changed as the mitophagy cutoff increased, so that higher mitophagy in Fig. 8 corresponds to fewer total mitochondria in the domain.

In Fig. 9, the value of *k*_*p*_ was also kept constant, but the resulting health levels were normalized by the total actual number of mitochondria in the domain. Because properly normalized health levels are not sensitive to the absolute number of mitochondria, this procedure was equivalent to explicitly fixing the total number of mitochondria. A direct comparison for the CoG model with explicitly fixed *M* is provided in Supplemental Material (S7 Fig).

Optimization of normalized health over all parameters (Fig. 10) was carried out by running simulations with 100 independent iterations each for a 10 × 10 grid of *N*_*s*_ and *f*_*s*_ values, for each mitophagy cutoff *ϕ*. The normalized average health and normalized last region health was computed for each set of parameter values, and the maximum of each for a given *ϕ* was found. The parameters that led to this maximum were then used to run another set of 10 identical simulations of 100 iterations each, to compute the standard deviation in the resulting normalized health. Error bars in Fig. 10 correspond to the standard error of the mean from these independent replicates.

## Acknowledgements

We thank Aidan Brown and Gulcin Pekkurnaz for helpful discussions.

## Funding Sources

Funding for this work was provided by the National Science Foundation CAREER Award (#1848057) and by the Hellman Fellows Fund.

## Supporting information

### Supporting Data

Data for generating Figures 2 through 9, as well as Matlab [83] scripts to generate those figures are provided as Supplementary Material.

### Generalization to Branched Axons

**Fig S1.**
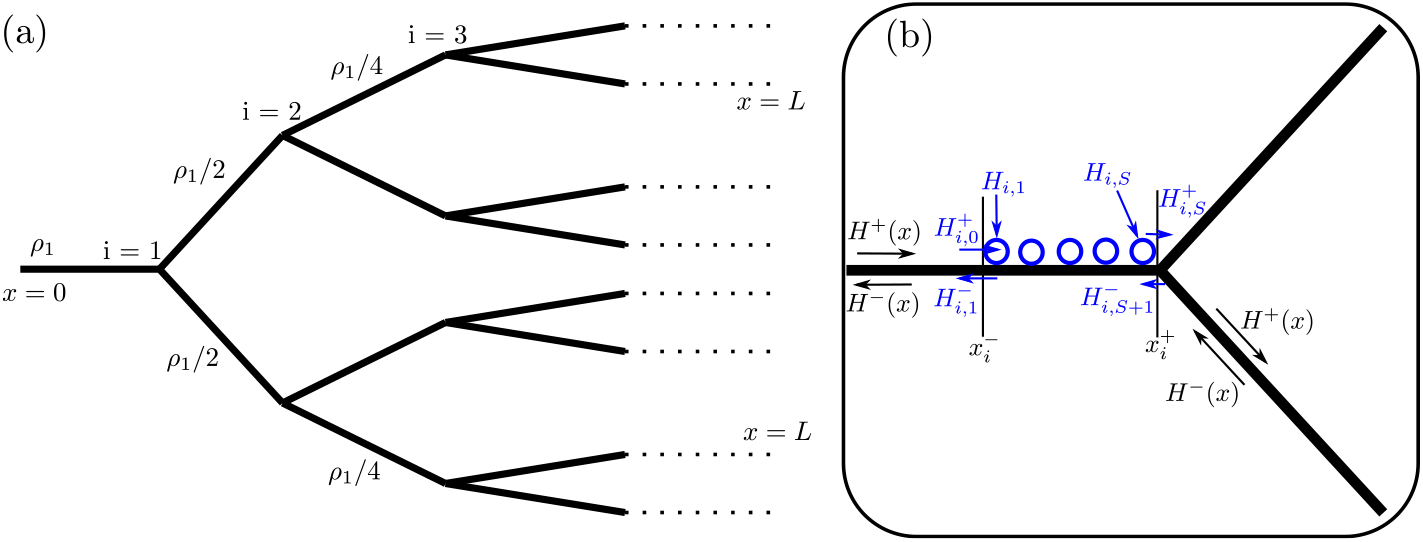
(a) Symmetric tree network used for branched axon calculations. Each segment branches into *g* = 2 identical downstream segments. The soma is at position *x* = 0 and distal tips at position *x* = *L*. (b) Zoomed-in schematic of a demand site at a branching junction. Demand site is located at position *x*_*i*_, with *S* = 5 stationary mitochondria shown (blue). The health of the *j*^th^ mitochondrion is given by *H*_*i,j*_, and the motile health leaving and entering on each side of the region is labeled. In our simplified model, the demand site is assumed to be infinitely narrow, and 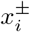 refers to the positions in the domain immediately after and before the demand site.

Throughout this work we focus on the interplay between mitochondrial transport, interchange, and aging, assuming a linear geometry to minimize the geometric complexity. However, *in vivo* axons have a tree-like branched structure, with mitochondrial localization observed at the branching points [29]. In this section we present a generalization of the ‘Space Station’ model for a simple symmetric tree structure with stationary mitochondria localized at the branching points.

We assume that each branching junction splits into *g* = 2 identical downstream branches of equal length. The “demand sites” are placed immediately upstream of each branch point, with an equal number of mitochondria (*S*) situated at each site. The model geometry is sketched in Fig. S1. The symmetry of the system allows us to define a coordinate system 0 ≤ *x* ≤ *L*, with 0 corresponding to the soma and *L* to the distal tips. The motile mitochondria health distribution 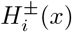 gives the health density at position *x* in any one of the corresponding branches. As before, we define *H*_*i,j*_ to be the health of the *j*^th^ mitochondrion at demand site (junction) *i*. The quantity 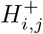 gives the anterograde-moving health density in the infinitesimally small space between mitochondrion *j* and *j* + 1 at site *i*; similarly, 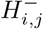 gives the retrograde-moving health density between mitochondrion *j* − 1 and *j*. With these definitions, the branched system obeys Equations 13 after replacing the general mitochondrial density *ρ* with a branch-dependent density *ρ*_*i*_, defined by

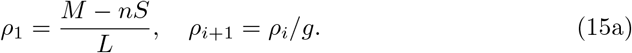

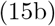

**Fig S2.**
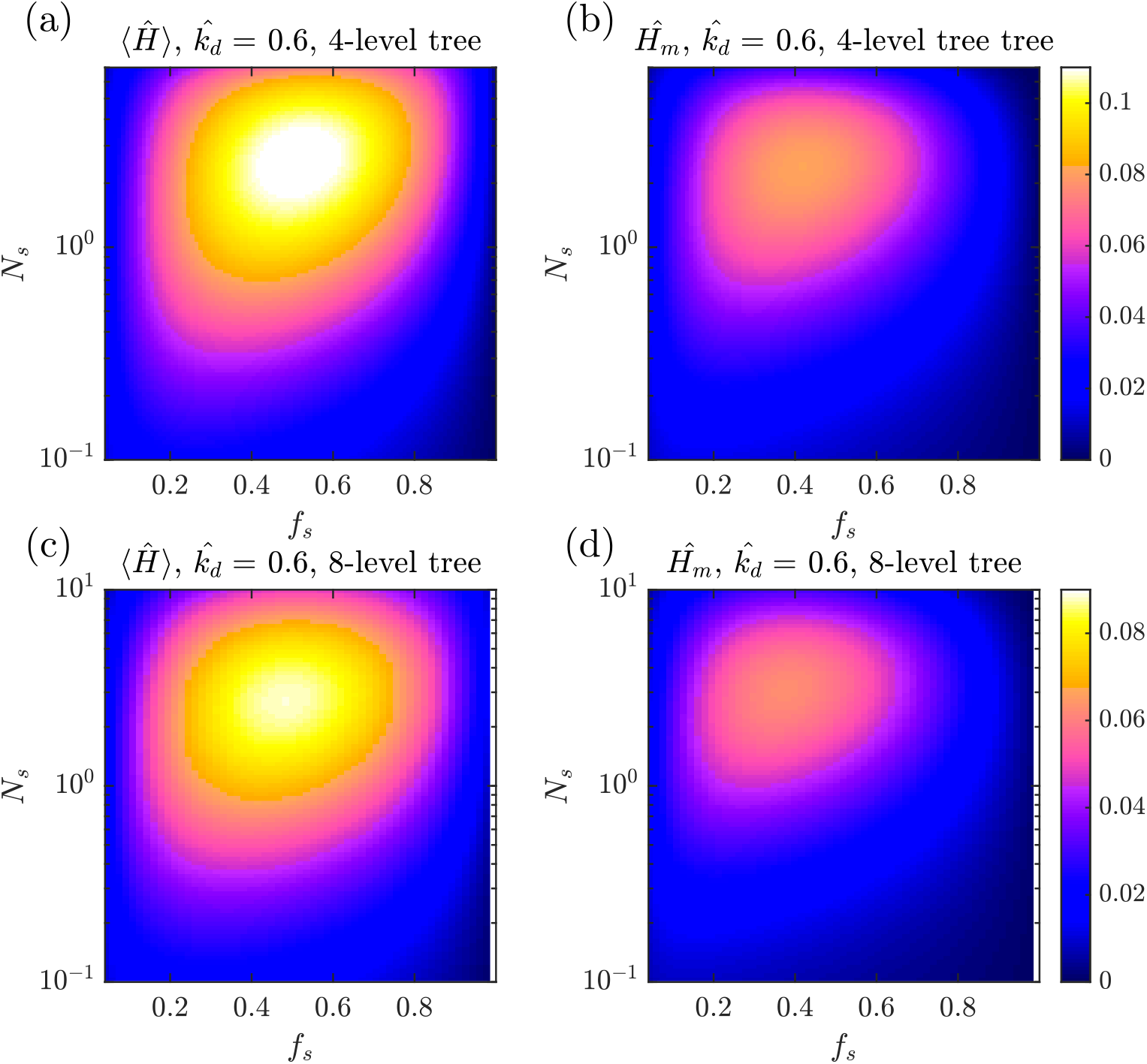
Mitochondrial health for expanded model in a symmetric branched tree geometry. (a) Average mitochondrial health as a function of *f*_*s*_ and *N*_*s*_ for a tree of depth *m* = 4, with *n* = 15 demand sites at the branch junctions. (b) Corresponding plot for the health of each distal demand site (furthest from the soma). (c-d) Corresponding plots for a tree of depth *m* = 8, with 255 demand sites.

Here *ρ*_1_ is the motile mitochondria density in the initial branch arising from the soma, and this density splits evenly at each junction point to give the downstream density *ρ*_*i*_ between junction *i* − 1 and *i*.

In addition to Eq. 13, the boundary conditions that complete the branched system are:

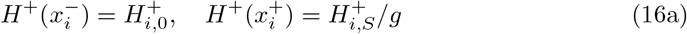

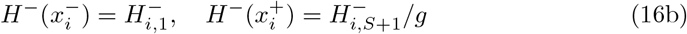

Eq. 16a indicates that the anterograde health density leaving each junction 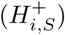 splits into *g* equal branches. Similarly, Eq. 16b defines the retrograde density entering the junction 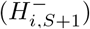 as the sum of retrograde densities from *g* branches.

The branched model with a tree of depth *m* has a total of *n* = 2^*m*^ − 1 demand sites, with each motile mitochondrion passing *m* of those sites on its way down the axon. When comparing to the linear model, we compare systems with the same total number of mitochondria *M* servicing the same number of demand sites *n*, and with the same distance *L* from soma to distal tip. It should be noted that the average linear density of motile mitochondria is lower in the branched model because the same total number *M* is spread out over a larger total branch length [*L*_tot_ = (2^(*m*+1)^ − 1)*L/*(*m* + 1)]. The primary model parameters (decay rate 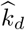, fraction of stopped mitochondria *f*_*s*_, and average number of stopping events for each protein *N*_*s*_) are defined to be conceptually analogous to the linear model. As before, we have 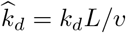 and *f*_*s*_ = *nS/M*. Because each mitochondrion traverses only one branch at each level of the tree, the number of stopping events is given by 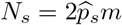.

The steady-state mitochondrial health at the demand sites in a tree of depth 4-level and 8-level tree are plotted in Fig. S2. The overall value of both average and distal mitochondrial health is somewhat decreased, presumably as a result of the lower density of motile mitochondria servicing the more distal branches. Interestingly, increasing the depth of the branching tree (while keeping a constant length *L*) only slightly lowers mitochondrial health, despite the fact that the distal density of motile mitochondria decreases exponentially. This result further confirms the observation that the primary relevant parameters are fraction of mitochondria stopped (*f*_*s*_) and number of stopping events *N*_*s*_ rather than the absolute number of demand sites or density of motile mitochondria. Furthermore, we note that the optimal values of *f*_*s*_ and *N*_*s*_ are largely unchanged in the branched system when compared to the linear geometry (Fig. 5). We therefore conclude that our main results, which rely on a linear axonal geometry, are more generally applicable.

A number of questions remain regarding mitochondrial maintenance in a branching geometry. Namely, the potential effect of redistributing stationary mitochondria at different depths along the tree, the consequences of asymmetric tree geometries, the effect of bidirectional motion into multiple branches, and the role of autophagy in tree-like structures, may further elucidate the optimal strategies for mitostasis in realistic axonal geometries. These more in-depth explorations serve as a promising jumping-off point for future expansion of the model described in this manuscript.

### Contribution of retrograde fusion events

Both the SS and CoG models allow health components to transition from a motile to a stationary state in an unbiased fashion. This means mitochonria moving in both the anterograde and retrograde directions are able to stop at demand sites (CoG model) or fuse with stationary mitochondria (SS) model with equivalent stopping or fusion probabilities. Because retrograde-moving mitochondria tend to have lower health levels compared to anterograde-moving mitochondria, removing fusion events between retrograde-moving mitochondria and stationed mitochondria is expected to result in higher health levels at demand sites. Such a modification would be equivalent to mitophagy that is triggered not by mitochondrial health levels but rather by the arrival of mitochondria at the distal terminus.

The ‘Space Station’ model equations can be modified as follows to reflect a potential regulatory mechanism that completely prohibits fusion of retrograde-moving mitochondria:

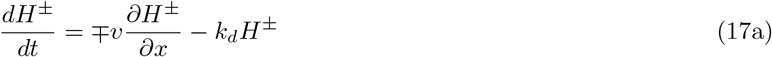

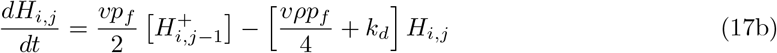

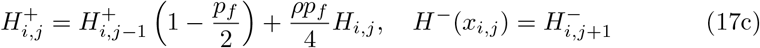

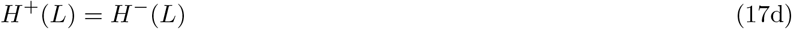

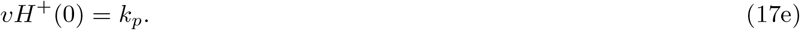

In Eq. 17b, the quantity *ρ/*2 gives the density of anterograde-moving mitochondria, replacing *ρ* in the original ‘Space Station’ equation (Eq. 13b).

**Fig S3.**
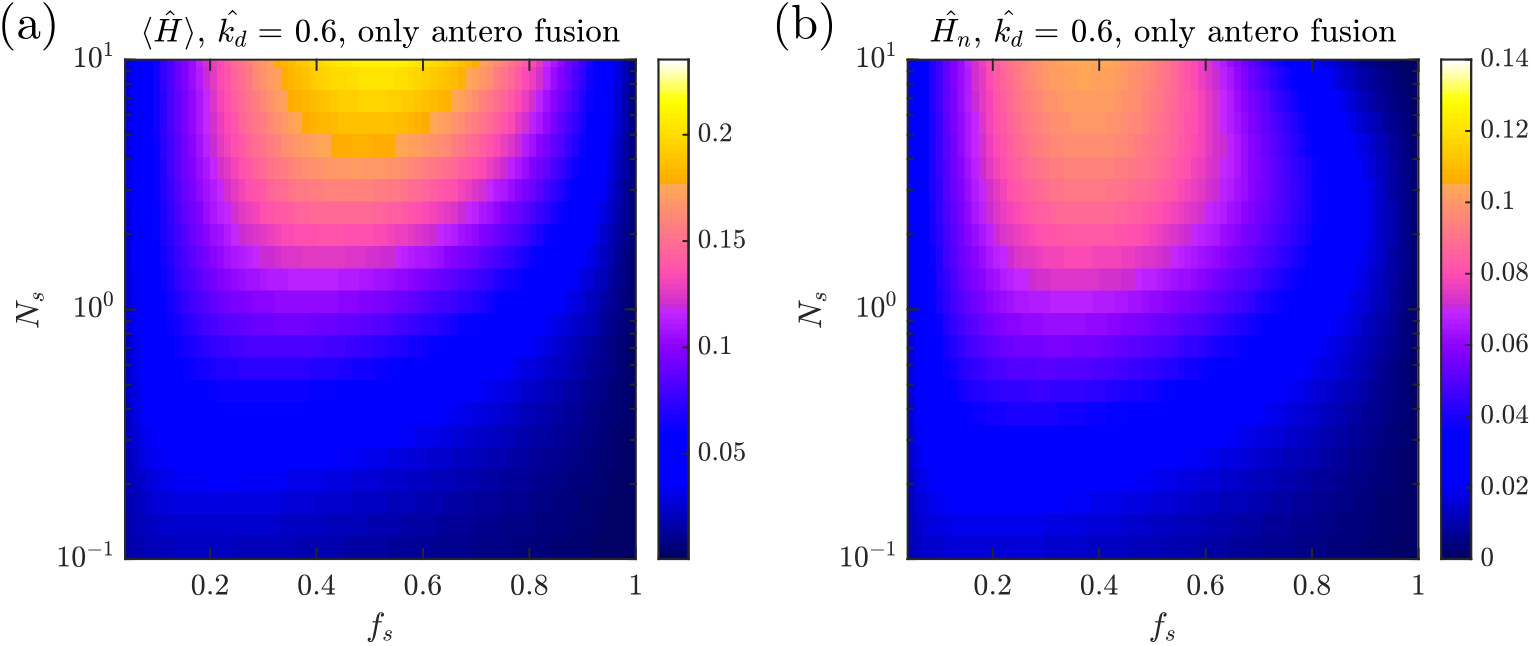
Mitochondrial health without retrograde fusion; plots analogous to Fig. 5. (a) Average health across all demand regions as a function of fraction of stopped mitochondria (*f*_*s*_) and number of stopping events (*N*_*s*_), for high decay rate 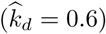. (b) Corresponding mitochondrial health at the most distal demand site.

The steady-state mitochondrial health in the absence of retrograde fusion is plotted in Fig. S3. We see that removing fusion with retrograde mitochondria means that there is no longer an optimum in the number of interaction events between motile and stationary mitochondria. The disadvantage to high *N*_*s*_ in the original model arose from the fusion of unhealthy retrograde mitochondria picking up proteins from stationary organelles and carrying them prematurely back to the soma for recycling. This disadvantage is no longer present if retrograde mitochondria are incapable of fusion.

Removal of retrograde fusion also results in a 50% increase in average health at demand sites (Figure S3a), again by preventing fusion of the less healthy retrograde mitochondria with stationary organelles at the proximal sites. Interestingly, the maximum health at the most distal site remains largely unchanged (Figure S3b).

These results indicate that any cellular mechanism capable of biasing exchange events between the motile and stationary population so that retrograde mitochondria were less likely to fuse or stop could be beneficial for enhancing mitochondrial health in the domain.

### Additional Supporting Figures

**Fig S4.**
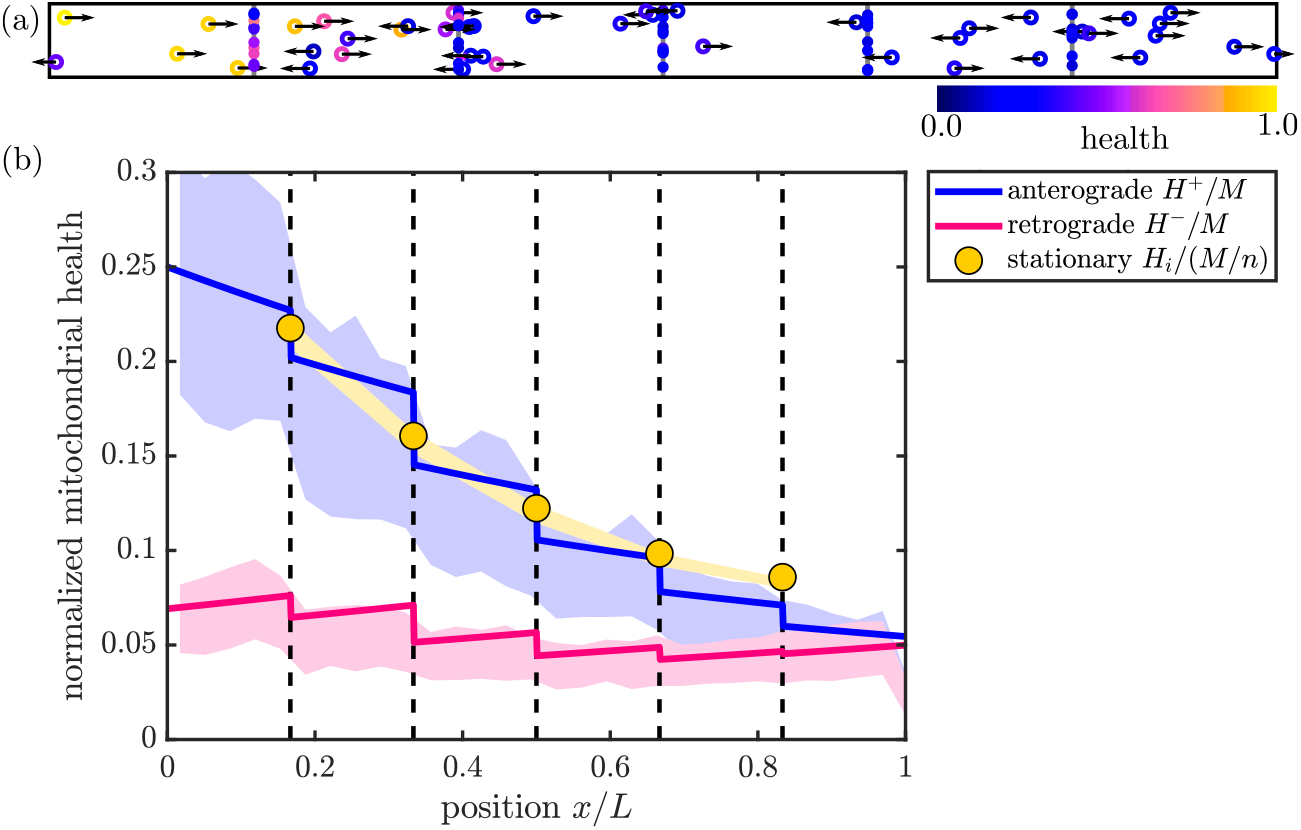
Figure supplement for Fig. 2. (a) Snapshot of a stochastic simulation for the SS model, with *M* = 100. (b) Steady-state solution for mitochondrial health in the SS model. Solid curves show linear density of mitochondrial health in anterograde (blue) and retrograde (magenta) mitochondria, normalized by total number of mitochondria in the domain. Yellow circles show total health at each of the discrete demand sites (dashed black lines), normalized by the total number of mitochondria per region. Shaded regions show corresponding quantities from discrete stochastic simulations (mean ± standard deviation) with *M* = 1500 mitochondria in the domain. Parameters used in (a) and (b): *n* = 5, *p*_*s*_ = 0.4, *f*_*s*_ = 0.5, *k*_*d*_*L/v* = 0.6.

**Fig S5.**
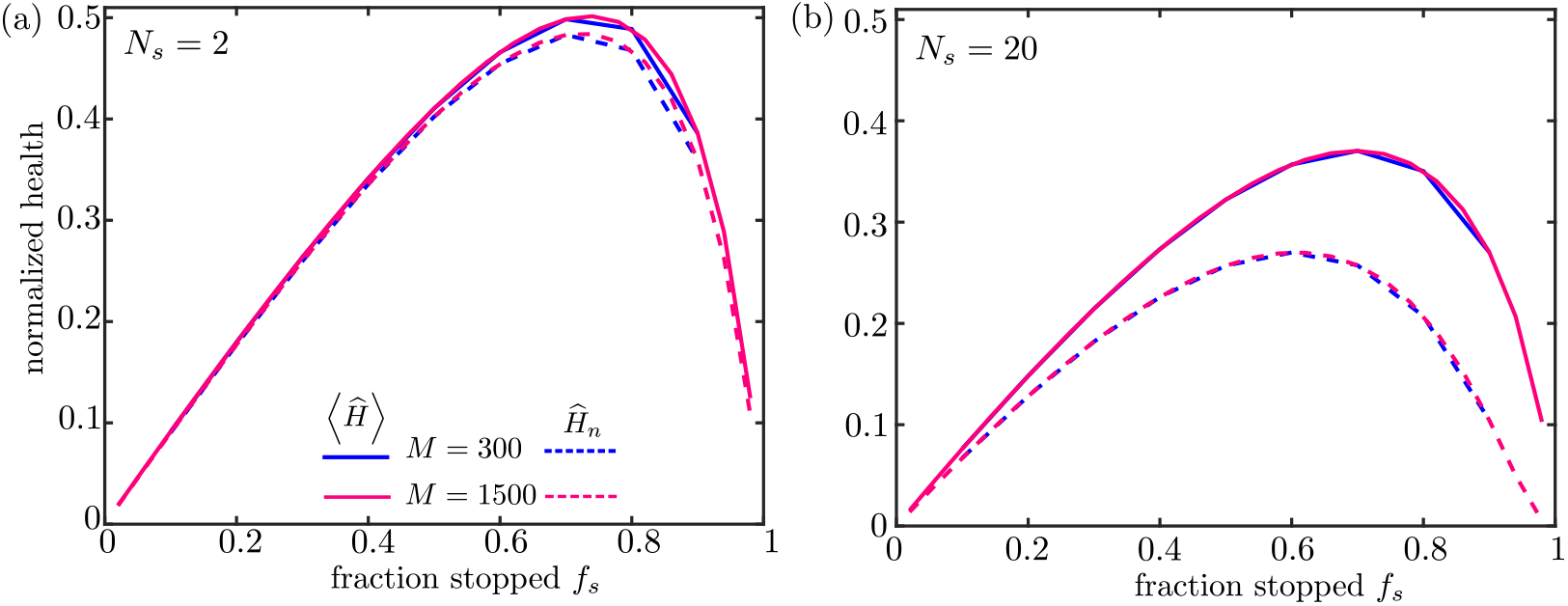
Figure supplement for Fig. 4. Mitochondrial health as a function of key dimensionless parameters. Solid curves show normalized average health over all demand sites; dashed curves show normalized health at the most distal site. The total number of mitochondria is set to *M* = 300 (blue) or *M* = 1500 (magenta). For each fraction of stationary mitochondria the fusion probability is adjusted to give a fixed number of stopping events for an individual protein traversing the domain: (a) *N*_*s*_ = 2 and (b) *N*_*s*_ = 20. All values shown are for the SS model, with *n* = 30 and 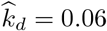.

**Fig S6.**
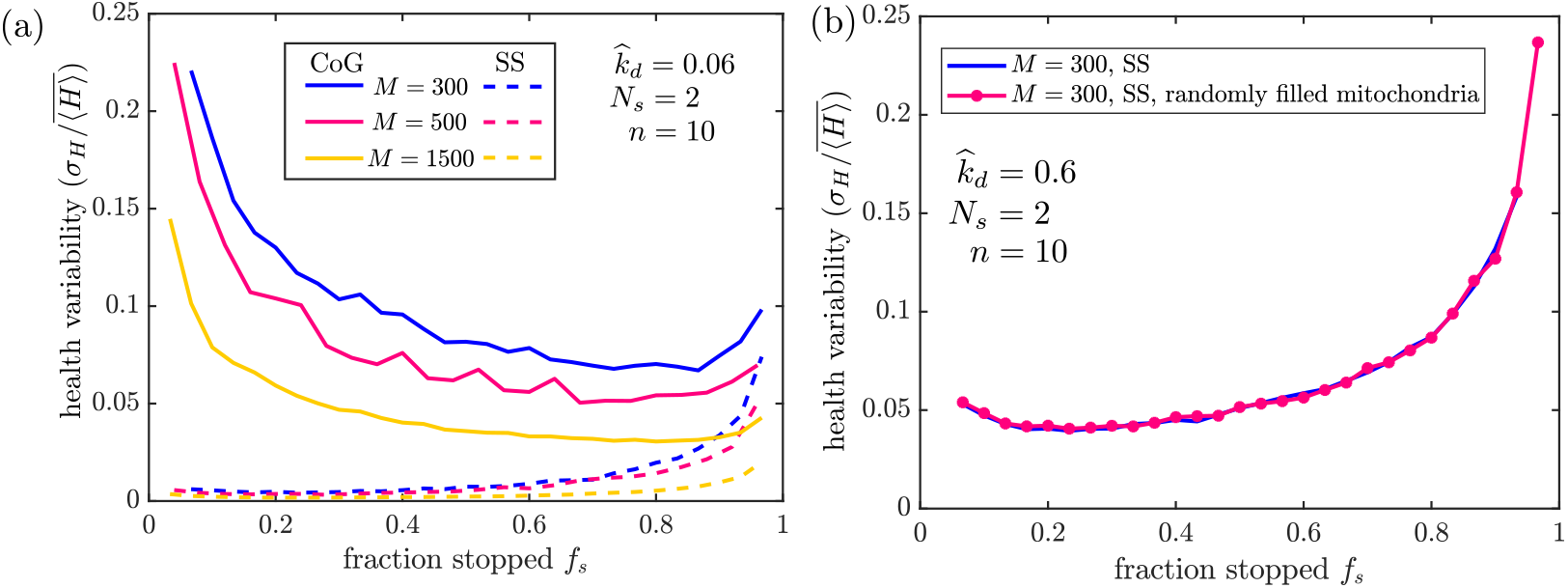
Figure supplement for Fig. 7. (a) Variability of mitochondrial health in different maintenance models, with low dimensionless decay rate 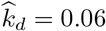. Results are shown for 300 (blue), 500 (magenta), and 1500 (yellow) average mitochondria in the domain. Solid lines correspond to simulations of the CoG model and dashed lines to the SS model. (b) Health variability in the SS model does not depend on how stationary mitochondria are distributed among sites. Results plotted are for decay rate 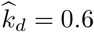, and number of mitochondria *M* = 300. Blue: equal number of mitochondria are placed at each demand site. Magenta: position of each stationary mitochondrion is selected uniformly at random among the demand site locations. Each iteration starts from an independently selected mitochondrial distribution. In all plots, variability is computed over 1000 indepedent iterations of stochastic simulations, using *n* = 10 demand sites and average stopping number *N*_*s*_ = 2.

**Fig S7.**
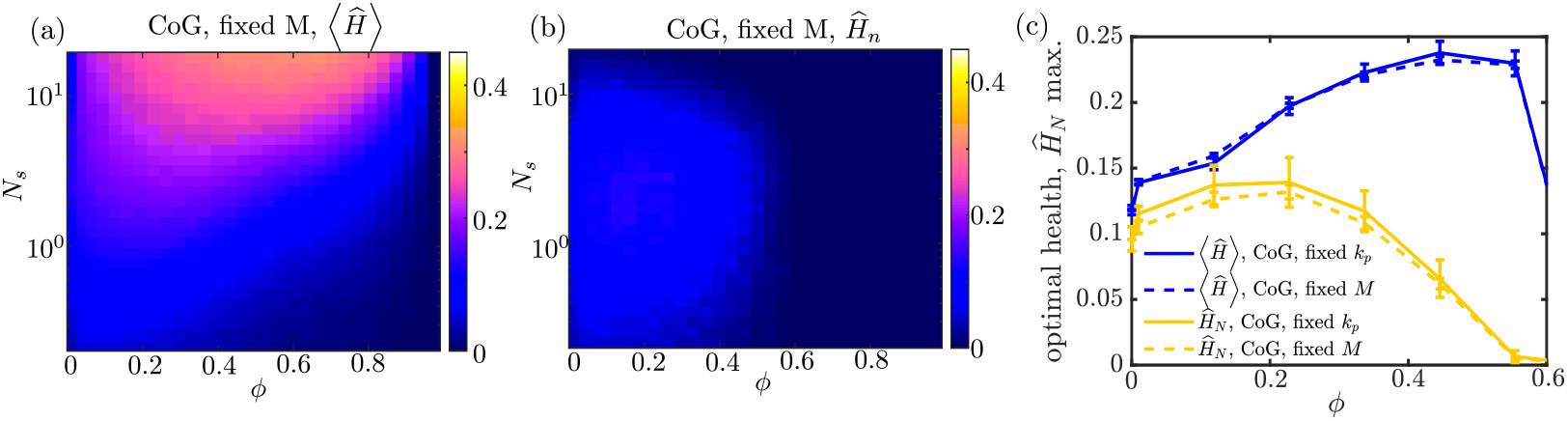
Figure supplement for Fig. 9. Rescaling health levels by total mitochondrial content is equivalent to a model with explicitly fixed number of mitochondria. (a-b) Average health across all regions (a), and health of last region (b), for simulations of a modified CoG model with fixed *M* = 150, to be compared to Fig. 9b-c. Health levels are normalized by *M/n* for both models. (c) Average health (blue) and last region health (yellow) for parameters that optimize the last region health at each fixed value of *ϕ*. Solid lines show the CoG model with *k*_*p*_ values fixed as the mitophagy threshold is varied (identical to curves in Fig. 10b). Dashed lines show corresponding plots for the model with explicitly fixed number of mitochondria *M*. Health levels normalized by *M/n* are shown to be largely the same in both cases.

**Fig S8.**
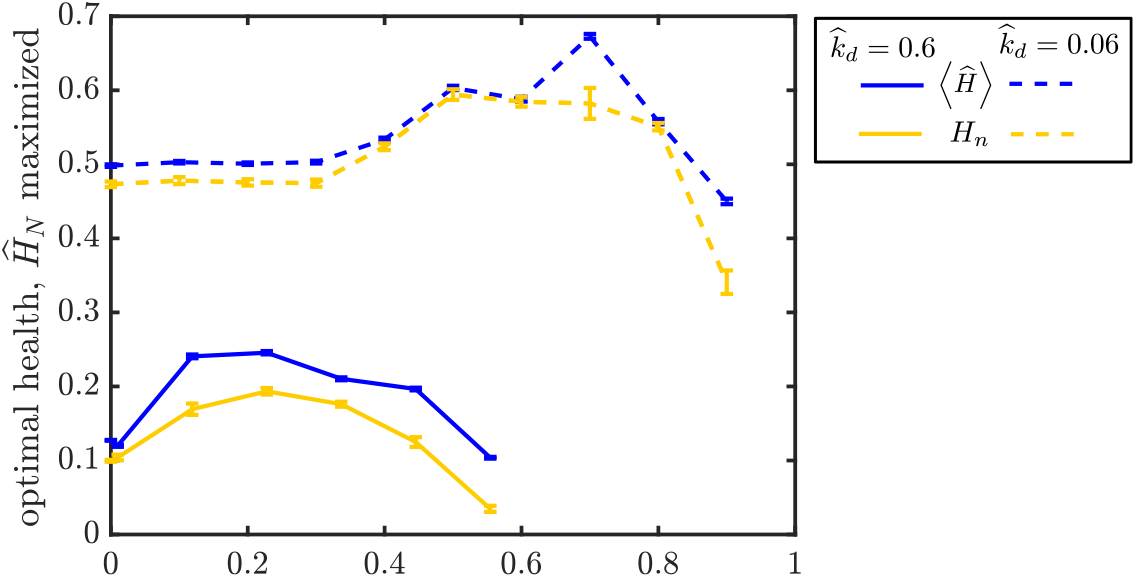
Figure supplement for Fig. 10. Optimal performance SS model in the presence of mitophagy, for different health decay rates. For each value of *ϕ*, the parameters *f*_*s*_ and *N*_*s*_ were optimized to yield the maximal health in the most distal region. Plotted are the resulting normalized health levels in the distal region (yellow) and averaged over all regions (blue). The health decay rate was set to 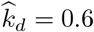 (solid) and 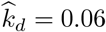 (dashed). Lower decay rates correspond to a higher mitophagy cutoff value *ϕ* that yields the maximal overall health. All health levels are normalized by total mitochondrial content per region (*M/n*), corresponding to a system where the total amount of mitochondrial material is limited. Error bars show standard error of the mean from 10 independent replicates.

**Fig S9.**
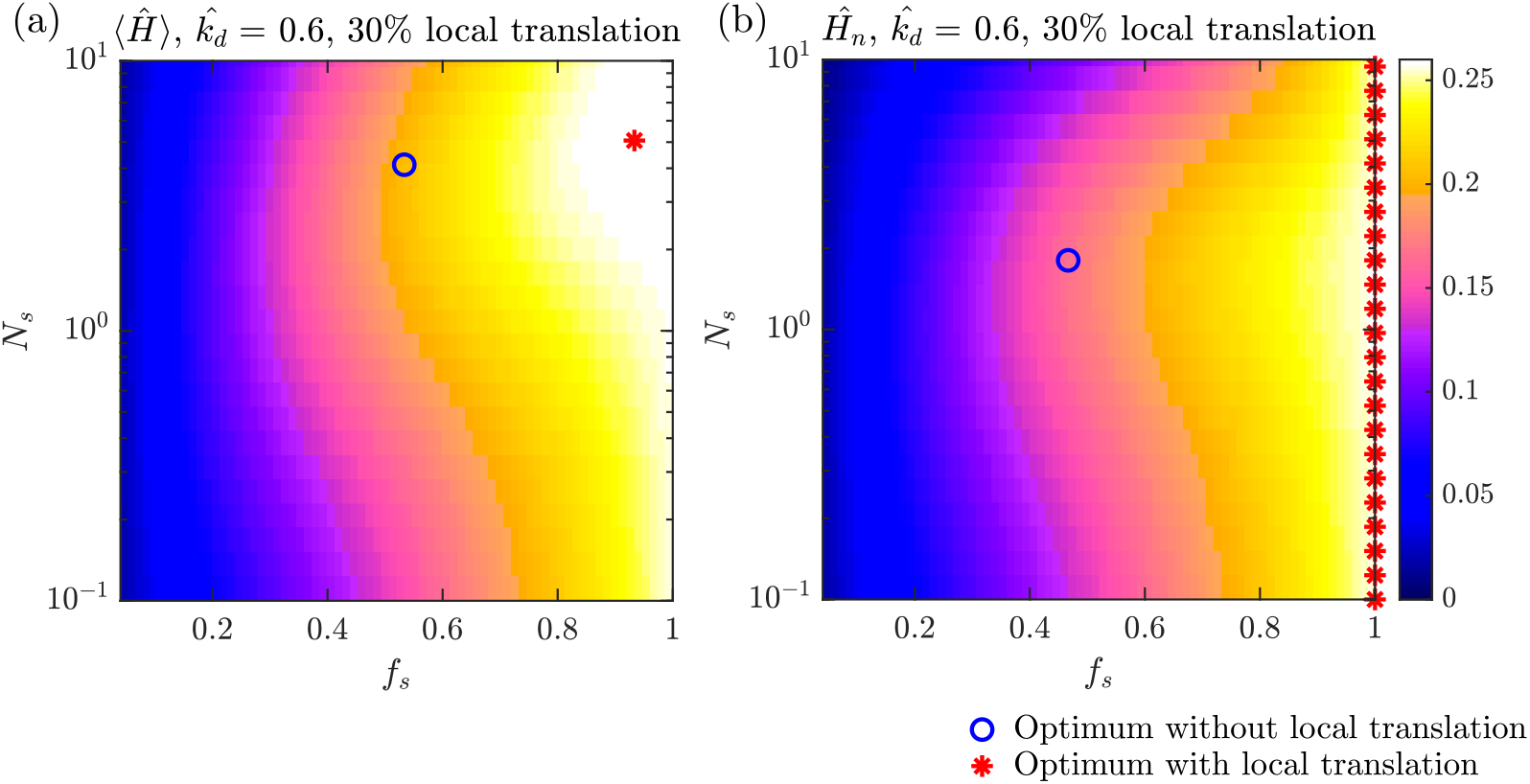
Figure supplement for Fig. 6. Mitochondrial health in the presence of substantial levels of local translation,with *α* = 30%. (a) Average health across all demand regions as a function of fraction of stopped mitochondria (*f*_*s*_) and number of stopping events (*N*_*s*_). Markers show parameters for optimal average health with local translation (red asterisk) and without it (blue open circle). (b) Corresponding plot for normalized mitochondrial health at the most distal demand site. Optimal distal health is obtained by placing all mitochondria in the stationary population (*f*_*s*_ = 1), so that health maintenance relies entirely on local translation rather than delivery from the soma.

## Notes

### Competing Interest Statement

The authors have declared no competing interest.

### Summary of Updates

Tree geometries and local translation included. See Agrawal and Koslover, PLoS Comp Biol, 2021 for final published version.

## References

1. González C, Couve A. The axonal endoplasmic reticulum and protein trafficking: Cellular bootlegging south of the soma. In: Seminars in cell & developmental biology. vol. 27. Elsevier; 2014. p. 23–31.

2. Lee SK, Hollenbeck PJ. Organization and translation of mRNA in sympathetic axons. Journal of Cell Science. 2003;116(21):4467–4478.

3. Kaplan BB, Gioio AE, Hillefors M, Aschrafi A. Axonal protein synthesis and the regulation of local mitochondrial function. In: Cell biology of the axon. Springer; 2009. p. 1–25.

4. Jung H, Yoon BC, Holt CE. Axonal mRNA localization and local protein synthesis in nervous system assembly, maintenance and repair. Nature Reviews Neuroscience. 2012;13(5):308–324.

5. Holt CE, Schuman EM. The central dogma decentralized: new perspectives on RNA function and local translation in neurons. Neuron. 2013;80(3):648–657.

6. Costa CJ, Willis DE. To the end of the line: axonal mRNA transport and local translation in health and neurodegenerative disease. Developmental neurobiology. 2018;78(3):209–220.

7. Sahoo PK, Smith DS, Perrone-Bizzozero N, Twiss JL. Axonal mRNA transport and translation at a glance. Journal of cell science. 2018;131(8).

8. Turner-Bridger B, Jakobs M, Muresan L, Wong HHW, Franze K, Harris WA, et al. Single-molecule analysis of endogenous β-actin mRNA trafficking reveals a mechanism for compartmentalized mRNA localization in axons. Proceedings of the National Academy of Sciences. 2018;115(41):E9697–E9706.

9. Zhou B, Yu P, Lin MY, Sun T, Chen Y, Sheng ZH. Facilitation of axon regeneration by enhancing mitochondrial transport and rescuing energy deficits. J Cell Biol. 2016;214(1):103–119.

10. Bradke F, Fawcett JW, Spira ME. Assembly of a new growth cone after axotomy: the precursor to axon regeneration. Nature Reviews Neuroscience. 2012;13(3):183–193.

11. Mar FM, Simões AR, Leite S, Morgado MM, Santos TE, Rodrigo IS, et al. CNS axons globally increase axonal transport after peripheral conditioning. Journal of Neuroscience. 2014;34(17):5965–5970.

12. Goldstein AY, Wang X, Schwarz TL. Axonal transport and the delivery of pre-synaptic components. Current opinion in neurobiology. 2008;18(5):495–503.

13. Lipton DM, Maeder CI, Shen K. Rapid assembly of presynaptic materials behind the growth cone in dopaminergic neurons is mediated by precise regulation of axonal transport. Cell reports. 2018;24(10):2709–2722.

14. Guedes-Dias P, Holzbaur EL. Axonal transport: Driving synaptic function. Science. 2019;366(6462):eaaw9997.

15. Doyle M, Kiebler MA. Mechanisms of dendritic mRNA transport and its role in synaptic tagging. The EMBO journal. 2011;30(17):3540–3552.

16. Zhao J, Fok AHK, Fan R, Kwan PY, Chan HL, Lo LHY, et al. Specific depletion of the motor protein KIF5B leads to deficits in dendritic transport, synaptic plasticity and memory. Elife. 2020;9:e53456.

17. Burow DA, Umeh-Garcia MC, True MB, Bakhaj CD, Ardell DH, Cleary MD. Dynamic regulation of mRNA decay during neural development. Neural development. 2015;10(1):11.

18. Dörrbaum AR, Kochen L, Langer JD, Schuman EM. Local and global influences on protein turnover in neurons and glia. Elife. 2018;7:e34202.

19. Harris JJ, Jolivet R, Attwell D. Synaptic energy use and supply. Neuron. 2012;75(5):762–777.

20. Rangaraju V, Calloway N, Ryan TA. Activity-driven local ATP synthesis is required for synaptic function. Cell. 2014;156(4):825–835.

21. Ibrahim M, Butt A, Berry M. Relationship between myelin sheath diameter and internodal length in axons of the anterior medullary velum of the adult rat. Journal of the neurological sciences. 1995;133(1-2):119–127.

22. Chiu SY. Matching mitochondria to metabolic needs at nodes of Ranvier. The Neuroscientist. 2011;17(4):343–350.

23. Rangaraju V, Lauterbach M, Schuman EM. Spatially stable mitochondrial compartments fuel local translation during plasticity. Cell. 2019;176(1-2):73–84.

24. Harris JJ, Attwell D. The energetics of CNS white matter. J Neurosci. 2012;32(1):356–371.

25. Sheng ZH. Mitochondrial trafficking and anchoring in neurons: new insight and implications. J Cell Biol. 2014;204(7):1087–1098.

26. Zhang CL, Ho PL, Kintner DB, Sun D, Chiu SY. Activity-dependent regulation of mitochondrial motility by calcium and Na/K-ATPase at nodes of Ranvier of myelinated nerves. Journal of Neuroscience. 2010;30(10):3555–3566.

27. Ohno N, Kidd GJ, Mahad D, Kiryu-Seo S, Avishai A, Komuro H, et al. Myelination and axonal electrical activity modulate the distribution and motility of mitochondria at CNS nodes of Ranvier. Journal of Neuroscience. 2011;31(20):7249–7258.

28. Davis AF, Clayton DA. In situ localization of mitochondrial DNA replication in intact mammalian cells. The Journal of cell biology. 1996;135(4):883–893.

29. Misgeld T, Schwarz TL. Mitostasis in neurons: maintaining mitochondria in an extended cellular architecture. Neuron. 2017;96(3):651–666.

30. Schwarz TL. Mitochondrial trafficking in neurons. Cold Spring Harbor perspectives in biology. 2013;5(6):a011304.

31. Wang X, Schwarz TL. The mechanism of Ca2+-dependent regulation of kinesin-mediated mitochondrial motility. Cell. 2009;136(1):163–174.

32. Pekkurnaz G, Trinidad JC, Wang X, Kong D, Schwarz TL. Glucose regulates mitochondrial motility via Milton modification by O-GlcNAc transferase. Cell. 2014;158(1):54–68.

33. Kang JS, Tian JH, Pan PY, Zald P, Li C, Deng C, et al. Docking of axonal mitochondria by syntaphilin controls their mobility and affects short-term facilitation. Cell. 2008;132(1):137–148.

34. Wong MY, Zhou C, Shakiryanova D, Lloyd TE, Deitcher DL, Levitan ES. Neuropeptide delivery to synapses by long-range vesicle circulation and sporadic capture. Cell. 2012;148(5):1029–1038.

35. Williams AH, O’donnell C, Sejnowski TJ, O’leary T. Dendritic trafficking faces physiologically critical speed-precision tradeoffs. Elife. 2016;5:e20556.

36. Nicholls DG. Mitochondrial membrane potential and aging. Aging cell. 2004;3(1):35–40.

37. Balaban RS, Nemoto S, Finkel T. Mitochondria, oxidants, and aging. cell. 2005;120(4):483–495.

38. Sheng ZH, Cai Q. Mitochondrial transport in neurons: impact on synaptic homeostasis and neurodegeneration. Nature Reviews Neuroscience. 2012;13(2):77–93.

39. Itoh K, Nakamura K, Iijima M, Sesaki H. Mitochondrial dynamics in neurodegeneration. Trends in cell biology. 2013;23(2):64–71.

40. Vincow ES, Merrihew G, Thomas RE, Shulman NJ, Beyer RP, MacCoss MJ, et al. The PINK1–Parkin pathway promotes both mitophagy and selective respiratory chain turnover in vivo. Proceedings of the National Academy of Sciences. 2013;110(16):6400–6405.

41. Cajigas IJ, Tushev G, Will TJ, tom Dieck S, Fuerst N, Schuman EM. The local transcriptome in the synaptic neuropil revealed by deep sequencing and high-resolution imaging. Neuron. 2012;74(3):453–466.

42. Westermann B. Mitochondrial fusion and fission in cell life and death. Nature reviews Molecular cell biology. 2010;11(12):872–884.

43. Wang S, Xiao W, Shan S, Jiang C, Chen M, Zhang Y, et al. Multi-patterned dynamics of mitochondrial fission and fusion in a living cell. PloS one. 2012;7(5):e19879.

44. Cagalinec M, Safiulina D, Liiv M, Liiv J, Choubey V, Wareski P, et al. Principles of the mitochondrial fusion and fission cycle in neurons. Journal of cell science. 2013;126(10):2187–2197.

45. Hoitzing H, Johnston IG, Jones NS. What is the function of mitochondrial networks? A theoretical assessment of hypotheses and proposal for future research. Bioessays. 2015;37(6):687–700.

46. Viana MP, Brown AI, Mueller IA, Goul C, Koslover EF, Rafelski SM. Mitochondrial fission and fusion dynamics generate efficient, robust, and evenly distributed network topologies in budding yeast cells. Cell systems. 2020;.

47. Liu X, Weaver D, Shirihai O, Hajnóczky G. Mitochondrial ‘kiss-and-run’: interplay between mitochondrial motility and fusion–fission dynamics. The EMBO journal. 2009;28(20):3074–3089.

48. Seager R, Lee L, Henley JM, Wilkinson KA. Mechanisms and roles of mitochondrial localisation and dynamics in neuronal function. Neuronal Signaling. 2020;4(2).

49. Youle RJ, Narendra DP. Mechanisms of mitophagy. Nature reviews Molecular cell biology. 2011;12(1):9–14.

50. Maday S, Wallace KE, Holzbaur EL. Autophagosomes initiate distally and mature during transport toward the cell soma in primary neurons. Journal of Cell Biology. 2012;196(4):407–417.

51. Mouli PK, Twig G, Shirihai OS. Frequency and selectivity of mitochondrial fusion are key to its quality maintenance function. Biophysical journal. 2009;96(9):3509–3518.

52. Patel PK, Shirihai O, Huang KC. Optimal dynamics for quality control in spatially distributed mitochondrial networks. PLoS Comput Biol. 2013;9(7):e1003108.

53. Figure generated with the aid of Servier Medical Art, licensed under a Creative Common Attribution 3.0 Generic License. http://smart.servier.com/.;.

54. Glock C, Heumüller M, Schuman EM. mRNA transport & local translation in neurons. Current opinion in neurobiology. 2017;45:169–177.

55. Yang E, van Nimwegen E, Zavolan M, Rajewsky N, Schroeder M, Magnasco M, et al. Decay rates of human mRNAs: correlation with functional characteristics and sequence attributes. Genome research. 2003;13(8):1863–1872.

56. Schwanhäusser B, Busse D, Li N, Dittmar G, Schuchhardt J, Wolf J, et al. Global quantification of mammalian gene expression control. Nature. 2011;473(7347):337–342.

57. Tushev G, Glock C, Heumüller M, Biever A, Jovanovic M, Schuman EM. Alternative 3/ UTRs modify the localization, regulatory potential, stability, and plasticity of mRNAs in neuronal compartments. Neuron. 2018;98(3):495–511.

58. Kim Y, Zheng X, Ansari Z, Bunnell MC, Herdy JR, Traxler L, et al. Mitochondrial aging defects emerge in directly reprogrammed human neurons due to their metabolic profile. Cell reports. 2018;23(9):2550–2558.

59. Wehnekamp F, Plucińska G, Thong R, Misgeld T, Lamb DC. Nanoresolution real-time 3D orbital tracking for studying mitochondrial trafficking in vertebrate axons in vivo. Elife. 2019;8:e46059.

60. Ashrafi G, Schlehe JS, LaVoie MJ, Schwarz TL. Mitophagy of damaged mitochondria occurs locally in distal neuronal axons and requires PINK1 and Parkin. Journal of Cell Biology. 2014;206(5):655–670.

61. Mandal A, Wong HTC, Pinter K, Mosqueda N, Beirl A, Lomash RM, et al. Retrograde mitochondrial transport is essential for organelle distribution and health in zebrafish neurons. Journal of Neuroscience. 2020;.

62. Miller KE, Sheetz MP. Axonal mitochondrial transport and potential are correlated. Journal of cell science. 2004;117(13):2791–2804.

63. Cai Q, Zakaria HM, Simone A, Sheng ZH. Spatial parkin translocation and degradation of damaged mitochondria via mitophagy in live cortical neurons. Current Biology. 2012;22(6):545–552.

64. Plucińska G, Paquet D, Hruscha A, Godinho L, Haass C, Schmid B, et al. In vivo imaging of disease-related mitochondrial dynamics in a vertebrate model system. Journal of Neuroscience. 2012;32(46):16203–16212.

65. Ferree AW, Trudeau K, Zik E, Benador IY, Twig G, Gottlieb RA, et al. MitoTimer probe reveals the impact of autophagy, fusion, and motility on subcellular distribution of young and old mitochondrial protein and on relative mitochondrial protein age. Autophagy. 2013;9(11):1887–1896.

66. Saxton WM, Hollenbeck PJ. The axonal transport of mitochondria. J Cell Sci. 2012;125(9):2095–2104.

67. Caminiti R, Carducci F, Piervincenzi C, Battaglia-Mayer A, Confalone G, Visco-Comandini F, et al. Diameter, length, speed, and conduction delay of callosal axons in macaque monkeys and humans: comparing data from histology and magnetic resonance imaging diffusion tractography. Journal of Neuroscience. 2013;33(36):14501–14511.

68. Fonkeu Y, Kraynyukova N, Hafner AS, Kochen L, Sartori F, Schuman EM, et al. How mRNA localization and protein synthesis sites influence dendritic protein distribution and dynamics. Neuron. 2019;103(6):1109–1122.

69. Wang X, Winter D, Ashrafi G, Schlehe J, Wong YL, Selkoe D, et al. PINK1 and Parkin target Miro for phosphorylation and degradation to arrest mitochondrial motility. Cell. 2011;147(4):893–906.

70. Twig G, Elorza A, Molina AJ, Mohamed H, Wikstrom JD, Walzer G, et al. Fission and selective fusion govern mitochondrial segregation and elimination by autophagy. The EMBO journal. 2008;27(2):433–446.

71. Zheng YR, Zhang XN, Chen Z. Mitochondrial transport serves as a mitochondrial quality control strategy in axons: Implications for central nervous system disorders. CNS Neuroscience & Therapeutics. 2019;25(7):876–886.

72. Magrané J, Sahawneh MA, Przedborski S, Estévez ÁG, Manfredi G. Mitochondrial dynamics and bioenergetic dysfunction is associated with synaptic alterations in mutant SOD1 motor neurons. Journal of Neuroscience. 2012;32(1):229–242.

73. Niescier RF, Kwak SK, Joo SH, Chang KT, Min KT. Dynamics of mitochondrial transport in axons. Frontiers in cellular neuroscience. 2016;10:123.

74. Lewis Jr TL, Turi GF, Kwon SK, Losonczy A, Polleux F. Progressive decrease of mitochondrial motility during maturation of cortical axons in vitro and in vivo. Current Biology. 2016;26(19):2602–2608.

75. Faits MC, Zhang C, Soto F, Kerschensteiner D. Dendritic mitochondria reach stable positions during circuit development. Elife. 2016;5:e11583.

76. Smit-Rigter L, Rajendran R, Silva CA, Spierenburg L, Groeneweg F, Ruimschotel EM, et al. Mitochondrial dynamics in visual cortex are limited in vivo and not affected by axonal structural plasticity. Current Biology. 2016;26(19):2609–2616.

77. Kubo R. The fluctuation-dissipation theorem. Reports on progress in physics. 1966;29(1):255.

78. Lehner B, Kaneko K. Fluctuation and response in biology. Cellular and Molecular Life Sciences. 2011;68(6):1005–1010.

79. Sato K, Ito Y, Yomo T, Kaneko K. On the relation between fluctuation and response in biological systems. Proceedings of the National Academy of Sciences. 2003;100(24):14086–14090.

80. Wolf L, Silander OK, van Nimwegen E. Expression noise facilitates the evolution of gene regulation. Elife. 2015;4:e05856.

81. Bodi Z, Farkas Z, Nevozhay D, Kalapis D, Lázár V, Csörgő B, et al. Phenotypic heterogeneity promotes adaptive evolution. PLoS biology. 2017;15(5):e2000644.

82. Yang Z, Steele DS. Effects of cytosolic ATP on spontaneous and triggered Ca2+-induced Ca2+ release in permeabilised rat ventricular myocytes. The Journal of Physiology. 2000;523(1):29–44.

83. MATLAB. version 9.5 (R2018b). Natick, Massachusetts: The MathWorks Inc.; 2018.

